# Ecotypes, *Wolbachia,* and urbanization shape *Culex pipiens* population structure in a West Nile virus hotspot

**DOI:** 10.64898/2026.05.10.722846

**Authors:** Sonia Cebrián-Camisón, Angel G. Rivera-Colón, Scott T. Small, Jordi Figuerola, Andrew D. Kern, Peter L. Ralph, Aureliano Bombarely, María José Ruiz-López

## Abstract

*Culex pipiens* is a major vector of West Nile Virus (WNV) in Europe and has a complex evolutionary history that has been linked to the probability of WNV spillover to humans. Here, we present a population genomic analysis of *Cx. pipiens* from a WNV hotspot in southwestern Spain. Whole-genome sequencing of 217 individuals from 24 localities revealed that population structure is mainly shaped by the coexistence of *pipiens* and *molestus* ecotypes. Despite clustering, genome-wide divergence between ecotypes was low, consistent with shared ancestral variation. Furthermore, *Wolbachia* infection was heterogeneous across the study area, where some *Culex pipiens pipiens* remain uninfected. We also identified three new polymorphic chromosomal inversions on chromosomes 1 and 3 that segregate independently. Although inversions did not underlie ecotypic differentiation, they were enriched for genes involved in olfaction and insecticide resistance pathways, suggesting potential adaptive and epidemiological relevance. Distance-based redundancy analyses showed that ecotype identity explained the largest fraction of genetic variance, with additional contributions from *Wolbachia* infection, habitat type (urban versus rural) and geographic distance. These results reveal that partial ecotypic differentiation, symbiont infection, and habitat interact to shape population structure in *Cx. pipiens*, with implications for vector ecology and disease transmission.

## Background

*Culex* mosquitoes are major disease vectors, transmitting pathogens affecting both animal and human health, including West Nile virus (WNV), the leading cause of viral encephalitis worldwide ^1^. Over the last two decades, WNV has spread widely across Europe, causing significant outbreaks in humans and horses ^2,3^. This recent increase in WNV incidence highlights the urgent need to better understand the ecological and evolutionary processes shaping *Culex* populations and their contribution to WNV transmission ^4^.

*Culex pipiens* is one of the primary vectors of WNV in Europe ^5^ and has a complex evolutionary history that has been linked to the probability of WNV spillover to humans ^6,7^. *Cx. pipiens s.s.* is one of the most widespread species within the *Cx. pipiens* complex, and it occurs across most temperate regions worldwide ^8^. Its broad distribution is likely facilitated by its tolerance to anthropized environments ^9^. This species comprises two morphologically indistinguishable ecotypes, *Cx. pipiens pipiens* and *Cx. pipiens molestus* (hereafter referred to as *pipiens* and *molestus*). These ecotypes have traditionally been considered to differ in key biological and behavioral traits, including reproductive strategy (autogeny vs. anautogeny), mating environment (confined vs. open spaces), feeding preferences (bird vs. mammal), and the occurrence of diapause. However, these traditional distinctions are largely generalizations based on comparisons between populations of *pipiens* living aboveground and populations of *molestus* living underground in northern Europe ^10^. In southern Europe this distinction is less clear. At lower latitudes, milder winters allow year-round persistence of both ecotypes ^11^, favoring aboveground co-occurrence of *pipiens* and *molestus* populations and potential hybridization ^12,13^. This has been linked to important epidemiological implications, because such hybrids may feed on both birds and humans, increasing WNV spillover ^14,15^. Recent whole-genome data have furthered our understanding of the evolutionary history of both ecotypes, and suggest that genetic similarity observed in southern populations reflects not only ongoing gene flow, but also ancestral variation predating the divergence of the form *molestus* ^16^. Capturing both fine-scale gene flow and complex evolutionary histories is therefore essential to understand pathogen transmission dynamics and inform control strategies.

A key genetic contributor to the formation and maintenance of ecotypes in mosquitoes is structural variation, particularly chromosomal inversions. Chromosomal inversions suppress recombination and maintain locally adaptive haplotypes under gene flow ^17,18^. In mosquitoes, chromosomal inversions have been found to influence traits linked to vectorial capacity such as biting and resting behavior ^19^, or susceptibility to *Plasmodium falciparum* in *Anopheles* ^20^. Although several inversions have been reported in the *Cx. pipiens* complex, e.g., in *Cx. quinquefasciatus* ^21^, these remain poorly characterized, limiting our understanding of their evolutionary significance and relevance for both vectorial capacity and ecotype divergence.

Beyond genomic architecture, the endosymbiont *Wolbachia* is an important factor influencing the vectorial capacity of mosquitoes as it can modify mosquito biology and their ability to transmit pathogens ^22,23^. In the case of *Cx. pipiens s.s., Wolbachia* seems to modulate WNV infection, but the evidence remains scarce and the directionality of this effect is unclear ^24^. In addition, *Wolbachia* can influence disease dynamics through its potential effect on population structure and gene flow caused by cytoplasmic incompatibility (CI). CI is a mechanism that prevents the development of offspring when sperm from *Wolbachia-*infected males fertilize uninfected eggs, or eggs infected by a non-compatible *Wolbachia* strain ^23^. This process may generate post-mating reproductive barriers that promote genetic divergence and can potentially drive speciation ^25,26^. In *Cx. pipiens s.s.*, infection with the *wPip* lineage is widespread but highly variable among regions and populations, with reported prevalence ranging from complete absence to near fixation and differences in *Wolbachia* strain composition ^27,28^. This heterogeneity creates a mosaic of reproductive compatibility among populations, with direct consequences for gene flow and population connectivity that have rarely been explored. In fact, *Wolbachia* CI may even be contributing to ecotype differentiation, but the extent to which this occurs remains unclear ^29,30^.

Vector population genetic structure and patterns of gene flow determine the spatial scale of pathogen transmission, dispersal potential and local vector competence, all of which critically affect disease epidemiology ^31^. Despite recent advances in the population genomics of *Cx. pipiens s.s.,* the relative contributions of ecological factors, *Wolbachia*-mediated reproductive barriers, and genomic architecture to population differentiation remain poorly understood. This knowledge gap is especially evident in regions with intense WNV transmission, where fine-scale population structure is likely to have direct epidemiological consequences.

Southern Spain has experienced endemic WNV circulation since the early 2000s ^32^, with large human outbreaks in 2020 and 2024 (77 and 158 human cases, respectively) ^33,34^. In addition, in this area, *Cx. pipiens* may play a key role as a bridge vector facilitating spillover of WNV to humans, particularly in urban environments ^35^. Previous studies also show that *Cx. p. pipiens and Cx. p. molestus* are present in southern Spain and show similar feeding patterns but different abundance in urban and rural regions, which might have implications for their respective roles in WNV amplification ^36^. This scenario, together with the recent evidence showing a complex evolutionary history of *Cx. Pipiens s.s.* in southern Europe ^16^, demonstrates that characterizing the genetic structure of *Cx. pipiens s.s.* populations in this region is essential. Here, we used population genomics to investigate how environmental factors and *Wolbachia* symbiosis interact to shape *Cx. pipiens s.s.* population structure in a WNV outbreak hotspot. Analyzing whole-genome data from 24 locations, we (1) characterized fine-scale population genomic structure and ecotype distribution, (2) quantified the relative contributions of habitat and *Wolbachia* infection to population differentiation, and (3) assessed the role of structural variants in these patterns. Our findings provide for the first time an integrated view of how ecological, symbiotic, and genomic factors jointly shape fine-scale mosquito population structure in a major WNV transmission hotspot, offering critical insights into WNV transmission dynamics and informing future vector control strategies.

## Results

### Genome-wide population structure of *Culex pipiens s.s.* is driven by ecotype differentiation

We sequenced 231 *Culex pipiens s.s.* individuals collected in 2022. Following molecular confirmation of species identity (Supplementary Figure 1), SNP calling, and quality filtering, 217 samples (213 females, 4 males) were included in the downstream analyses. These samples were collected from 24 trapping locations (7-10 individuals per site; except Barbate, n=2; Supplementary Table 1). In total, 4,535,359 SNPs were identified across the three chromosomes (range: 1.0-1.9 million SNPs per chromosome), with a mean read depth of 12.76x (s.d=1.17). After linkage disequilibrium (LD) pruning, we retained 596,248 SNPS for chromosome 1; 1,299,163 for chromosome 2 and 968,399 for chromosome 3. These were used to assess population structure using genome-wide Principal Component Analysis (PCA) and ADMIXTURE.

The genome-wide PCA showed a U-shaped pattern, with the first principal component (PC1) explaining 17.04% of the variance (Figure 1A). Samples were distributed along PC1 between two opposing ends connected by a continuous gradient, with most individuals occupying intermediate positions. A subset of individuals clustered at both extremes of PC1 (12 and 10 individuals, respectively) and displayed PC2 values over 0.1 (Figure 1A). ADMIXTURE analyses showed a pattern consistent with the PCA. The optimal number of genetic clusters was K=2, but most individuals (n=186) exhibited mixed ancestry (Figure 1C). Individuals with ancestry proportions exceeding 95% for either of the clusters corresponded to those located at opposite extremes of PC1 and showing higher PC2 values (Figure 1A).

**Figure 1:**
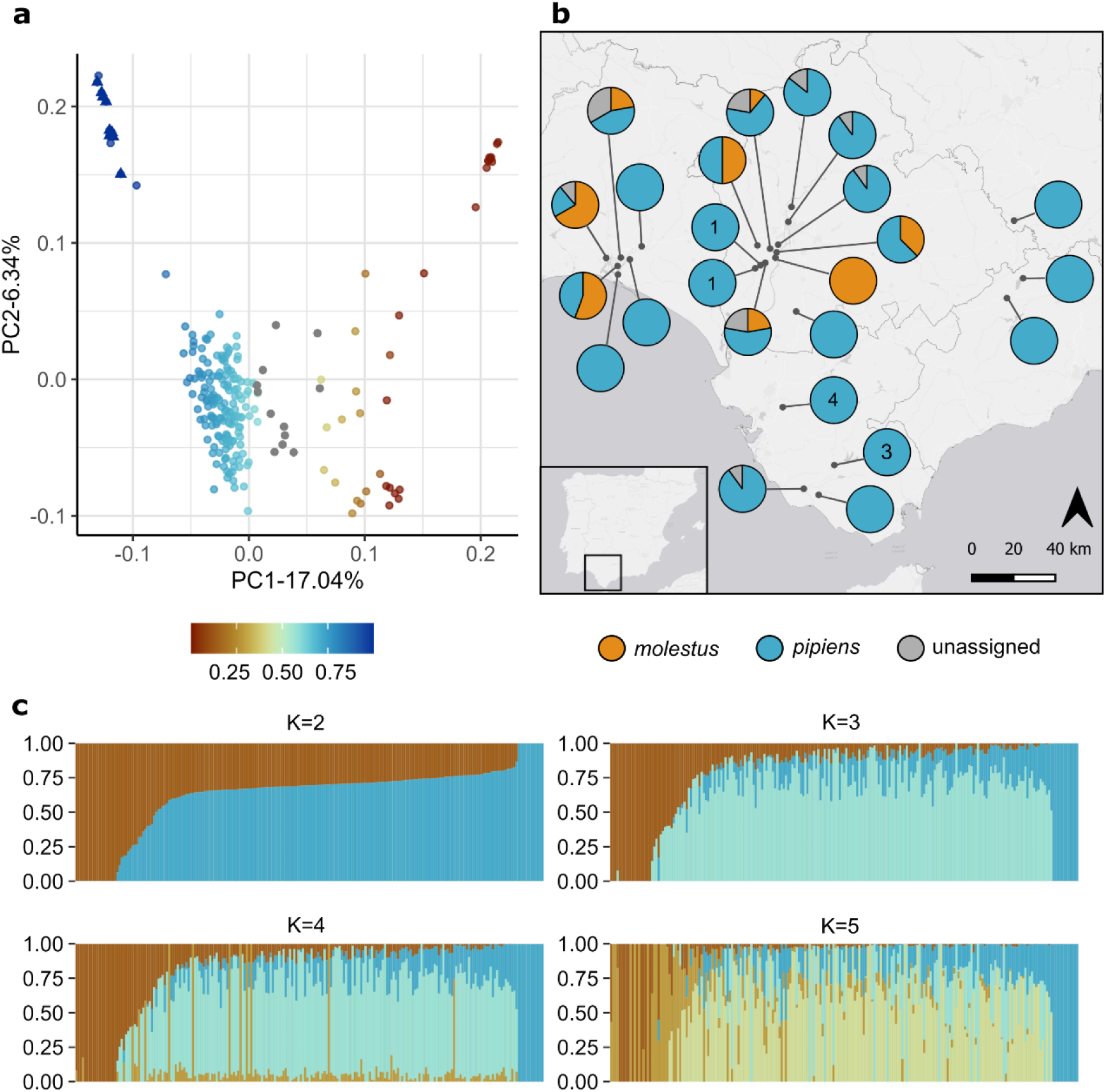
Population structure of *Culex pipiens s.s* in southwestern Spain. a) Principal Components Analysis (PCA) of 217 *Cx. pipiens s.s.* Color gradient represents estimated proportion of *Cx. p. pipiens* ancestry (p(Q2) in ADMIXTURE analyses with K=2): samples with higher *Cx. p. pipiens* ancestry are shown in blue, whereas those with higher *Cx. p. molestus* ancestry are shown in orange. Grey dots represent samples that could not be confidently assigned to any of the ecotypes. Triangles represent samples uninfected by *Wolbachia*. b) Map of sampling sites showing the proportion of *pipiens* and *molestus* samples among mosquitoes analyzed for each locality. Unassigned individuals are represented in grey. The number of mosquitoes not infected by *Wolbachia* is shown inside pie charts. c) ADMIXTURE results for K=2 to K=5, illustrating individual ancestry proportions. Individuals are ordered according to their estimated ancestry proportions rather than sampling locality.

To assess whether the observed population structure reflected ecotype differentiation, we ran a PCA on a combined dataset including our samples and type specimens with assigned ecotypes from ^16^ (Supplementary Table 2). In this analysis, *pipiens* and *molestus* reference specimens clustered at opposite ends of PC1, and our samples aligned along the same axis according to their ADMIXTURE ancestry proportions (Supplementary Figure 2). Based on their position relative to reference samples and ancestry proportions, we assigned individuals to ecotypes: 171 grouped with *Cx. p. pipiens* and 34 grouped with *Cx. p. molestus*. Twelve individuals showed intermediate ecotype ancestry (40-60%) and occupied intermediate positions along PC1, including regions where *pipiens* and *molestus* reference samples partially overlap. Because we could not assign these individuals to either ecotype based on ancestry and PCA position, they were left unassigned.

While PC1 primarily captures ecotype differentiation, PC2 revealed additional structure within ecotypes (Figure 1A). Within *pipiens*, PC2 separated a small group of individuals with ≥95% *pipiens* ancestry (n=12). Within *molestus*, PC2 separated specimens from Puebla del Río from all other populations. This site was the only sampling location where all individuals were assigned to *molestus* (Figure 1b) and showed >95% *molestus* ancestry. To confirm this result was not a consequence of relatedness we ran a PCA accounting for kinship coefficients. Individuals showed low relatedness with a mean kinship coefficient of 0.022 and this PCA was consistent with our main analysis (Supplementary Figure 3). A total of 54 samples with kinship coefficients ≥ 0.125 were excluded from the PCA. None of the excluded samples were from Puebla del Rio.

### Presence of polymorphic chromosomal inversions

We next examined how localized genomic divergence contributes to genome-wide patterns of variation. Windowed PCA (100 kb windows) revealed substantial genetic differentiation in three regions on chromosomes 1 and 3, separating samples in two or three distinct clusters (Figure 2B). These regions also showed high linkage disequilibrium (LD), with R^2^ values close to 1 (Figure 2A). PCA restricted to each region revealed three differentiated clusters along PC1 (Supplementary Figure 4). These LD and PCA patterns are consistent with chromosomal inversions. Each cluster formed in the PCA was identified as either the wildtype (i.e., arrangement whose SNPs were more similar to the reference genome), inverted, or the heterokaryotype. All three regions showed elevated F_ST_ relative to the genomic background (Figure 2C), with substantially higher differentiation when comparing the putative wildtype and inverted homokaryotypes, further supporting the presence of chromosomal inversions and the use of the PCA patterns as a proxy for karyotype.

**Figure 2:**
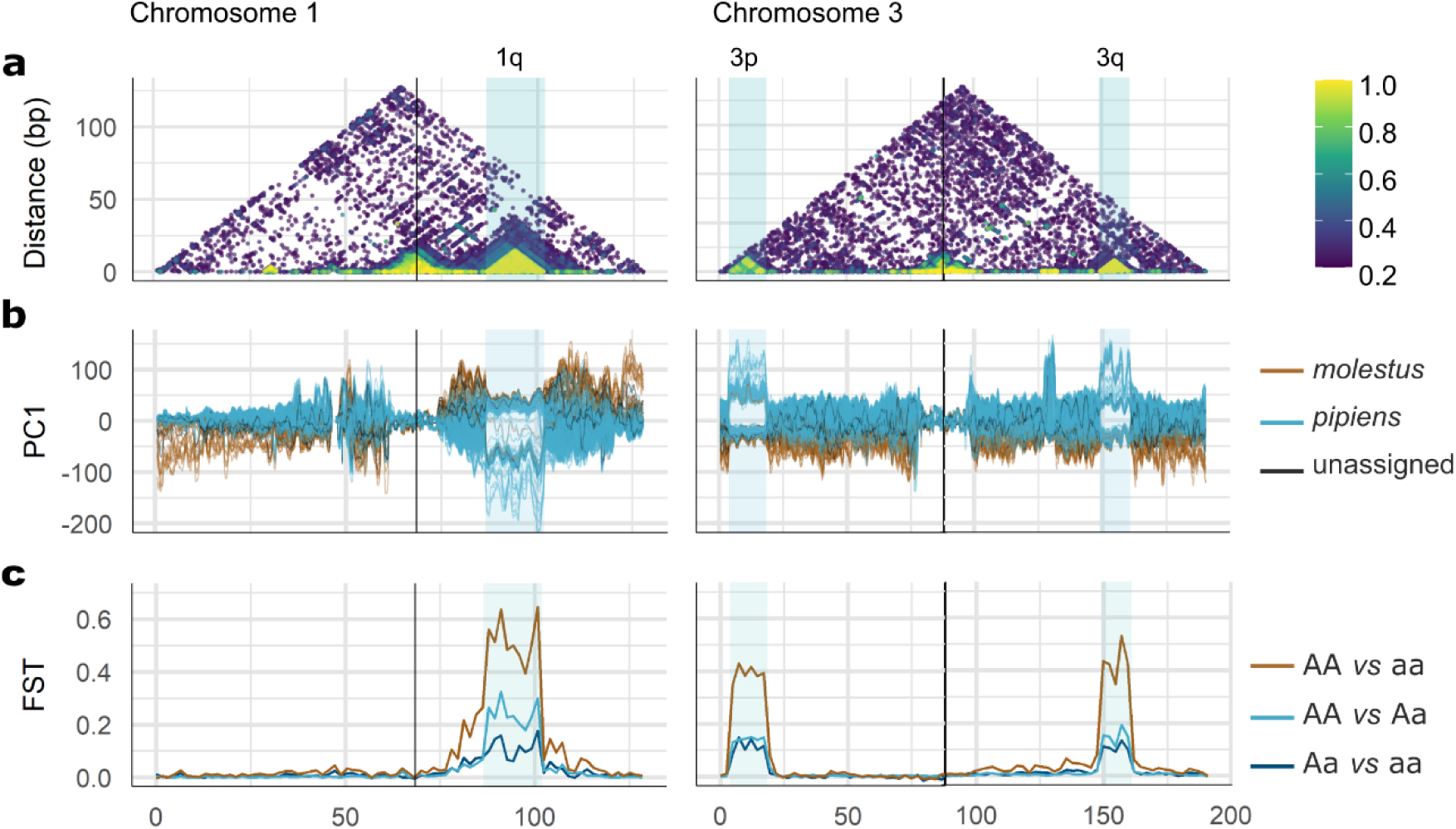
Genetic differentiations and linkage disequilibrium on chromosomal inversions from *Culex pipiens*. a) Linkage disequilibrium estimates (R^2^), b) winPCA results over 100 kb windows and c) F_ST_ estimates along chromosomes 1 and 3. Centromeres are indicated with vertical black lines. Shaded blue areas mark putative chromosomal inversions.

Candidate inversions were named following standard nomenclature used in *Anopheles*, based on the chromosomal location. On chromosome 1, the candidate inversion 1q spanned from approximately 87 to 102 Mb and contained 479 genes. On chromosome 3, two regions were identified: 3p (spanning from 3.8 to 18.5 Mb, containing 487 genes) and 3q (spanning from 149 to 161 Mb, containing 450 genes). Two genes were affected by the distal breakpoint of 1q and the proximal breakpoint of 3q. The frequency of the inverted arrangement was 17.9% for 1q, 16.8% for 3p and 74.4% for 3q. Although the three inversions were polymorphic in *pipiens*, for inversion 3q all *molestus* specimens carried the inverted homokaryotype. For 1q and 3p, 32 out of 34 *molestus* specimens carried the wildtype homokaryotype (Supplementary Table 3). Furthermore, karyotype assignment differed among the three inversions, indicating they are not genetically linked and segregate independently (Supplementary Figure 4).

GO enrichment analyses revealed significant functional enrichment for all three inversions after FDR correction (α=0.05; Supplementary Table 4). Inversion 3p showed the highest number of significantly enriched GO terms. These terms included numerous odorant receptors (GO:0050911, GO:0050896, GO:0004984), as well as cytochrome P450 genes (GO:0004497, GO:0020037, GO:0005506, GO:0016705) and ABC-type transporters (GO:0140359). Inversion 3q showed enrichment of genes like cuticle proteins (GO: 0042302) and alpha crystallins (GO:0005212).

Finally, to test whether inversion polymorphisms influenced the observed population genomic structure, we repeated the PCA and admixture analyses using a biallelic SNPs from chromosome arm 2q, which does not contain candidate inversions. This dataset retained 54,954 sites across 90 individuals with no missing genotypes. PCA and ADMIXTURE analyses yielded patterns consistent with those obtained using the three chromosomes (Supplementary Figure 5). The PCA remained primarily structured along PC1 by ecotypes, and ADMIXTURE supported K=2. These results confirm that the observed population structure was not confounded by fixed inversions karyotypes or missing genotypes.

### Low genome-wide divergence and incomplete genetic isolation between ecotypes

We quantified absolute (D_XY_ and D_A_) and relative (F_ST_) divergence between *pipiens* and *molestus* using 10kb genomic windows across the genome. Analyses were performed on a reduced “reference” dataset of 90 representative samples (see Methods), including biallelic SNPs and invariant sites without missing genotypes across all three chromosomes. This dataset matches the samples that were used on the PCA and ADMIXTURE analyses accounting for the effect of inversions and missing data. Values of F_ST_ and D_A_ were generally low across the whole genome (D_A_: mean=7.87·10^-5^, sd=1.53·10^-5^; F_ST_: mean=0.051, sd=0.072; Figure 3) while D_XY_ estimates revealed moderate divergence (mean=1.19·10^-3^, sd=1.22·10^-3^). Overall, these patterns show low levels of differentiation and divergence between ecotypes, which is consistent with recent gene flow and/or shared ancestry. Genome-wide nucleotide diversity (π) was similar in *pipiens* and *molestus* (*pipiens*: mean=1.19·10^-3^, sd=1.21·10^-3^; *molestus*: mean=1.02·10^-3^, sd=1.26·10^-3^). However, within chromosomal inversions, π was lower for *molestus* than for *pipiens* (Figure 3).

**Figure 3:**
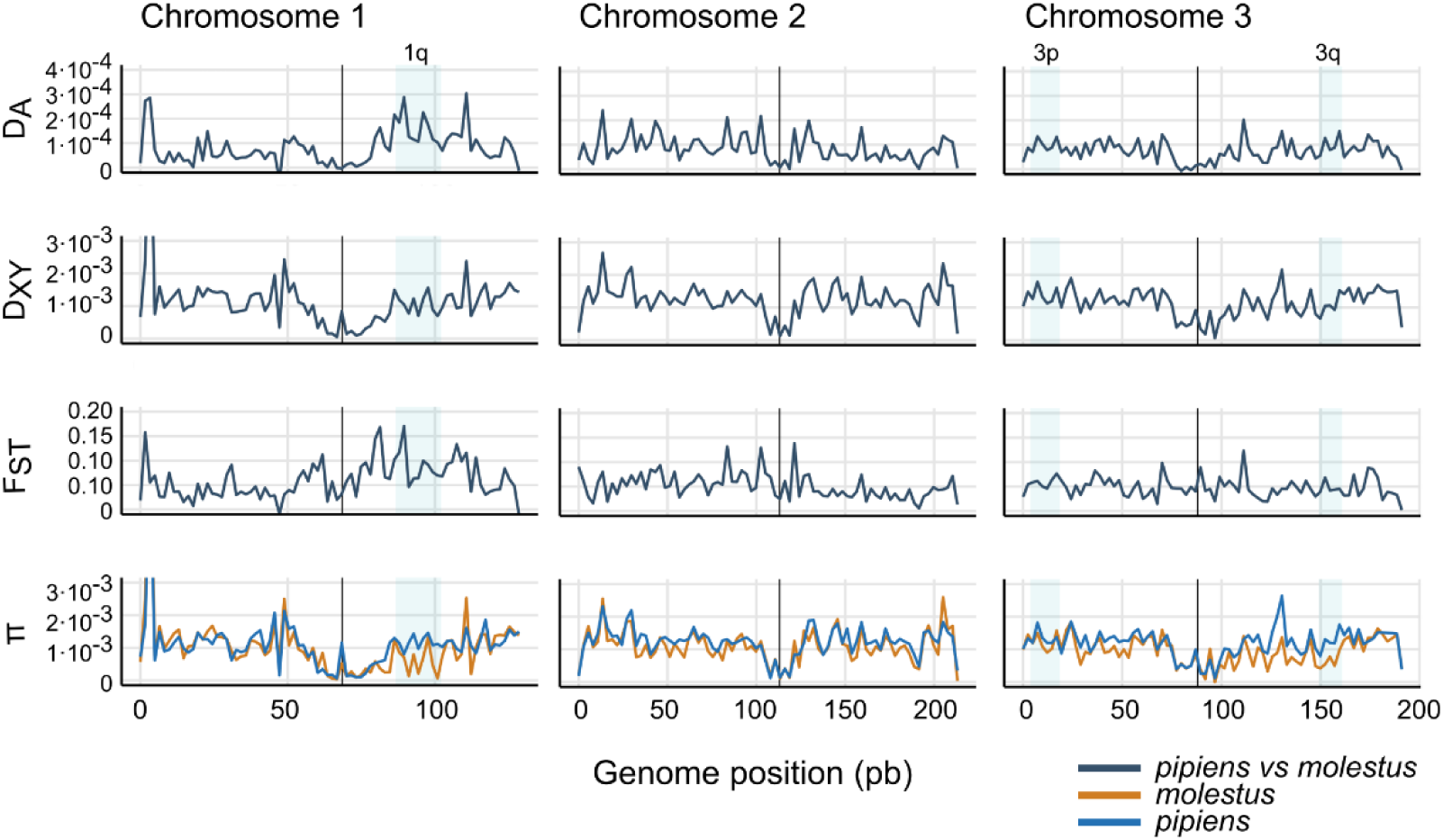
D_XY_, F_ST_ and DA between *pipiens* and *molestus* (ecotype ancestry >60%) and nucleotide diversity (π) estimated over 10kb genomic windows. Centromeres are indicated with vertical black lines. Shade blue areas mark putative chromosomal inversions.

When calculations were restricted to individuals with >95% ecotype ancestry, we observed high peaks of F_ST_ and D_A_ co-localizing with chromosomal inversions 1q and 3q (Supplementary Figure 6). At these regions, F_ST_ showed values up to 0.4 (1q) and 0.3 (3q), while D_A_ reached 0.001 (1q) and 7.5·10^-4^ (3q). No such increase was observed at inversion 3p. However, when we compared individuals sharing the same homokaryotype across ecotypes at 1q and 3q, F_ST_ values remained comparable to genome-wide background levels (Supplementary Figure 7). In contrast, we observed a substantial increase in F_ST_ at inversion 3p when comparing different homokaryotypes, regardless of whether they belonged to the same or different ecotypes. These results suggest divergence between ecotypes in inverted regions is primarily driven by the polymorphic state of these inversions, with different karyotypes becoming fixed among individuals with ancestry proportions above 95% (Supplementary Table 3). Moreover, these patterns of divergence further suggest that the origin of the inversions predates the split between ecotypes.

To further assess genetic relationships, we performed phylogenetic reconstruction using biallelic SNPs from the 90 reference samples, restricting the analyses to both arms of chromosome 2 to avoid any biases from inversions. Despite low overall differentiation of ecotypes, phylogenetic reconstruction clustered *pipiens* and *molestus* as two sister clades (Supplementary Figure 8). However, six individuals changed their clade assignment between chromosomal arms. Four had mixed ancestry and were unassigned, while the remaining two were one *molestus* and one *pipiens* with less than 95% ecotype ancestry. Notably, individuals with >95% *pipiens* ancestry consistently formed a monophyletic group, whereas individuals with >95% *molestus* ancestry did not. This phylogenetic pattern and the instability across chromosomal regions suggest a scenario of incomplete genetic isolation with some level of gene flow. Consistent with this hypothesis, *f_3_* statistics on biallelic SNPS from chromosome 2 detected significant admixture signals in *pipiens* (*f_3_* = -5.83E-04, Z = -3.46), whereas no such signal was observed in *molestus* (Supplementary Table 5). The unassigned group showed weak or no significant evidence of admixture between both ecotypes (|Z| ≤ 3), supporting their classification as unassigned.

To test whether ecotype divergence was driven by specific sets of genes, we carried out GO enrichment analyses based on 5kb and 10kb windows comparing the most divergent regions between *pipiens* and *molestus* (1% higher values of F_ST_ between ecotypes). However, no GO terms were significantly enriched, regardless of whether ecotypes were defined using >60% ancestry or >95% ancestry thresholds.

### *Wolbachia* infection prevalence varies with ecotype ancestry

After competitive mapping, we found that 118 out of the 127 (92.91%) *Cx. pipiens s.s.* individuals were infected by *Wolbachia.* Among infected individuals, 115 showed almost complete coverage of the *w*Pip genome (>98%), with a mean depth of 55.86X (range:6.03X-211.97X). The remaining three individuals had lower *w*Pip genome coverage (3.89, 4.71 and 46.66%, respectively) suggesting low infection intensity. All nine uninfected individuals, as well as the three with low coverage, were *Cx. p. pipiens* with ≥95% *pipiens* ancestry (Figure 1b). To identify *w*Pip groups (*w*Pip-I to *w*Pip-V) we performed BLAST searches against a curated panel of *pk1* reference sequences (Supplementary Table 6). All consensus assemblies matched the *w*Pip-I reference sequence (accession number OM885005.1)^37^, with full-length alignments and 100% identity. This indicated that all infected mosquitoes carried the *w*Pip-I strain and that the *pk1* marker was invariant in our samples. Consequently, no phylogenetic reconstruction was performed for this marker.

### Symbiotic, environmental and spatial factors influence ecotype-driven population structure

Consistent with our previous results, multivariate analyses confirmed that ecotype is the primary driver of genetic structure, while also revealing significant contributions from additional ecological and spatial factors. Distance-based redundancy analyses (dbRDA) explained 11.35% of the total genetic variance and the overall model was statistically significant (F = 3.33, p = 0.0001, df= 8). Ecotype was the strongest predictor, accounting for 50.81% of the constrained variation (F=13.53, p=1·10^-4^, df=1), while *Wolbachia* infection, habitat (urban/rural) and geographic structure (PCNMs derived from Euclidean distances among sampling locations) also contributed significantly (Table 1). Mantel tests also revealed significant correlations between geographic and genetic distances within each ecotype. However, the correlation was higher in *molestus* (r= 0.454; p=0.0001) than in *pipiens* (r=0.188, p=0.0001), indicating a much stronger pattern of isolation-by-distance in *molestus* populations. Beta-mixed effects model also revealed that mosquitoes collected in urban environments had significantly higher *molestus* ancestry than those from natural habitats (pQ1; β = 1.42 ± 0.59 SE, z = 2.42, p = 0.015).

**Table 1:** Summary of distance-based redundancy analyses (dbRDA). Results show the marginal effects of each predictor on genome-wide genetic distances among *Culex* samples. Significance levels are based on 9,999 permutations.

| Variable | Df | F | % Total<br>Variance | % Constrained<br>Variance | p-value |
| --- | --- | --- | --- | --- | --- |
| **Ecotype** | 1 | 13.53 | 5.76% | 50.81% | 0.0001*** |
| **Geography** | 5 | 1.11 | 2.36% | 20.79% | 0.039* |
| **Wolbachia infection** | 1 | 2.26 | 0.96% | 8.51% | 0.0002*** |
| **Habitat** | 1 | 1.88 | 0.80% | 7.06% | 0.0004*** |

## Discussion

Understanding how ecological and spatial factors, symbiont infection, and genomic architecture interact to shape mosquito population structure is essential for predicting vector-borne disease risk. Using whole-genome data from a West Nile virus (WNV) hotspot in southwestern Spain, we show that *Culex pipiens s.s.* fine-scale genomic population structure was primarily driven by differentiation between *pipiens* and *molestus* ecotypes, while maintaining extensive shared ancestry. This structure was further influenced by habitat, geographic distance and *Wolbachia* infection. In addition, we identified three polymorphic chromosomal inversions shared by both ecotypes. This pattern highlights the complex interactions between evolutionary and ecological factors that influence mosquito population connectivity and can ultimately drive disease transmission.

As expected, genome-wide PCA and ADMIXTURE analyses identified two genetic clusters corresponding to *pipiens* and *molestus*, yet most individuals exhibited mixed ancestry. This pattern is characteristic of Mediterranean populations, where both ecotypes frequently coexist aboveground ^12,13^. Early studies based on microsatellites ^13^ or single diagnostic loci ^36^, interpreted this pattern as evidence of ongoing hybridization. However, recent work using whole-genome data has shifted this interpretation. Haba *et al.*^16^ showed that much of the admixture observed in southern populations reflects the persistence of ancestral polymorphism predating the emergence of the *molestus* ecotype, rather than extensive contemporary hybridization. They also showed that historical gene flow mainly occurred from northern *molestus* populations into *pipiens*.

Our results are consistent with this pattern, with phylogenetic analyses separating *molestus* and *pipiens* ecotypes. Furthermore, individuals with intermediate ancestry classified as unassigned did not show clear evidence of recent admixture based on *f_3_* statistics and cannot be considered true hybrids. Instead, we found evidence of introgression from *molestus* into *pipiens,* which has also been observed in *Cx. pipiens s.s.* populations in Portugal ^13^. The small number of unassigned individuals reduces the power of *f_3_* statistics to detect ongoing gene flow. However, the combination of low genome-wide differentiation and within-ecotype structure, suggests that much of the apparent recent admixture corresponds to shared ancestry and past introgression. Still, ongoing gene flow cannot be completely dismissed. This updated genomic interpretation also helps reconcile earlier work based on the CQ11 marker ^36^, which likely captures a mixture of retained ancestral polymorphism and true introgression, thereby inflating estimates of contemporary hybridization ^16^.

Alongside ecotype, *Wolbachia* infection, geography, and habitat each explained small but significant proportions of the genetic variation. Similarly to other European populations ^29,30^, *Wolbachia* prevalence was over 90%. Notably, all the individuals classified as *pipiens* with >95% ancestry were either uninfected or showed low infection intensity likely acquired through recent horizontal acquisition ^23,38^. Although the potential contribution of distinct *Wolbachia* lineages to the reproductive isolation of *pipiens* and *molestus* ecotypes has been discussed ^29,30^, we did not find evidence supporting this hypothesis because all infections correspond to the same *w*Pip-I lineage. Thus, population structure may result from differences in infection frequency among populations, rather than from strain incompatibility. However, the observed infection pattern could also be caused by the presence of migrants originating from uninfected populations in our sampling locations. In this case, the associations between infection status and genetic structure would reflect recent demographic mixing rather than persistent CI. Either scenario could explain why individuals with >95% *pipiens* ancestry were separated along PC2 from other *pipiens*. Both possibilities also highlight the importance of accounting for symbiont dynamics when interpreting population structure, particularly in regions where *Wolbachia* prevalence is below 100%.

Habitat also contributed significantly to population structure, with mosquitoes from urban localities showing higher *molestus* ancestry than those from natural habitats. This supports previous observations that ecotype abundance varies along urbanization gradients in southern Spain ^36^. Furthermore, ecotype formation is often driven by genetic variants that predate the formation of contemporary ecotype pairs, persisting as standing variation or entering populations through introgressive hybridization ^39^.Such standing genetic variation provides a readily available substrate for rapid divergence when ecological conditions change. Our findings regarding urbanization reinforce that differentiation between *pipiens* and *molestus* largely reflects the differential sorting and reuse of ancient polymorphisms, rather than *de novo* mutation or extensive contemporary hybridization. This aligns with Haba *et al.* ^16^, who showed that *molestus* did not evolve as a genomic response to urbanization, but instead exploited ancestral standing variation to adapt to novel urban environments. Mantel tests showed stronger isolation-by-distance in *molestus* than in *pipiens*, suggesting more limited dispersal or stronger spatial structuring in *molestus*. This difference may reflect contrasts in breeding site stability, dispersal behavior, demographic history, or greater dependence on urban environments, consistent with their habitat associations.

We found no evidence of functional enrichment associated with ecotype differentiation, indicating that the genetic basis of ecotypic divergence is not strongly linked to specific genes. Instead, differentiation may involve subtle regulatory changes or polygenic variation, particularly in environments where ecological contrasts are weak. Thus, the limited genomic and functional differentiation observed in our populations may reflect the relatively similar ecological conditions experienced by *pipiens* and *molestus* in Mediterranean aboveground habitats. Overlapping breeding environments and comparable climatic regimes are likely to reduce the strength of divergent selection. By contrast, stronger ecological contrasts in northern regions may promote more pronounced differentiation in both genomic composition and gene expression, potentially explaining the greater ecotypic separation.

Although chromosomal inversions are considered a major driver of ecotype differences ^40^, the three chromosomal inversions identified here likely play a limited role on ecotype differentiation. Instead, they likely represent ancestral genetic polymorphism segregating in both ecotypes. While we were unable to obtain precise breakpoint estimates using software benchmarked to work with short-read data, the differentiation shown by the windowed PCA, the increased genetic divergence and the high linkage disequilibrium revealed in those regions are indirect but strong evidence for the presence of chromosomal inversions. Future cytogenetic analyses or long-read whole-genome resequencing would help to confirm the presence and resolve accurate breakpoint positions. Such experiments would also identify the genes affected by inversion breakpoints. Despite not contributing to ecotype differentiation, these inversions also carry adaptive or epidemiologically relevant variation. For example, we showed that inversion 3p is enriched for odorant receptors (ORs). Olfaction is strongly linked with important mosquito behaviors such as sugar-feeding, host-seeking, and oviposition ^41–43^. In addition, differential expression of odorant receptors is sometimes linked to insecticide resistance ^44,45^. More directly relevant to insecticide resistance, we demonstrated that 3p is enriched for cytochrome P450 genes and ABC-type transporters, genes involved in detoxification with well-stablished roles in metabolic resistance ^45–47^. Thus, inversions may harbor wide adaptive potential for *Cx. pipiens s.s.* within this region, especially in *pipiens,* which displayed a higher level of polymorphism. Similarly, inversion 3q, which appears differentially fixed across >95% ancestry individuals, is enriched in gene families commonly associated to resistance mechanisms, like cuticular proteins and alpha-crystallins, implicated in cuticular and stress-related resistance pathways, respectively ^44,45,48,49^. Further work is required to determine whether alternative inversion arrangements are associated with phenotypic differences in resistance levels.

It is often assumed that the *pipiens* and *molestus* ecotypes show preferential feeding on birds and mammals, respectively, and that hybrids show intermediate behavior ^50,51^. This idea supports the hypothesis that hybrids may act as bridge vectors, facilitating spillover of pathogens such as WNV and Usutu virus (USUV) to humans. However, evidence from southern Europe does not fully support this scenario. In the same area studied here, Martínez-de la Puente *et al*. ^36^ found no significant association between mosquito ecotype and blood-meal origin. Both ecotypes and their hybrids fed predominantly on birds, while a substantial proportion of blood meals (≈33%) were derived from mammals, including humans. Similar feeding patterns were reported in Portugal^52^, despite contrasting evidence ^51^. Our results provide an evolutionary context for this behavioral plasticity. The extensive shared ancestry, low genome-wide divergence and lack of ecotype-specific functional differences detected suggest that feeding behavior is unlikely to be fixed by ecotype in this region. Instead, host choice may be influenced by other factors such as prey availability, habitat type, or season ^53^. Puebla del Río illustrates this dynamic. This town was the epicenter of the 2020 and 2024 WNV outbreaks in Spain, where *molestus* was particularly abundant and viral circulation was high within the urban area ^35^. These findings demonstrate that intense transmission can occur even in places where one form dominates, without needing hybrids to explain the link between bird and human infection. In this situation, the role of hybrids may have been overstated. If *pipiens*, *molestus*, and intermediate genotypes all feed on both birds and mammals under favorable conditions, then any of them could contribute to spillover. In such a scenario, the diversity and local structuring of vector populations, combined with ecological opportunity, become determinants of epidemiological risk.

These findings also have practical implications for control and surveillance. Approaches that treat *Cx. pipiens* as a single, uniform population may miss key local differences. In urban areas, in particular, it is essential to consider the higher prevalence and spatial clustering of *molestus*, as well as to target below-ground habitats where *molestus* can persist and reproduce throughout the winter ^11^. At the same time, incorporating genomic information into surveillance and extending it to other villages and cities with strong WNV incidence, could help identify local differences that affect how populations respond to insecticides, *Wolbachia*-based strategies, or habitat management. This becomes essential in systems where *Cx. pipiens s.s.* form a genetic continuum rather than discrete evolutionary units. Such populations may harbor great adaptive potential that could contribute to buffer environmental changes through recombination and redistribution of adaptive variation, with great implications for disease transmission in the current global change context.

In conclusion, *Cx. pipiens s.s.* populations in southwestern Spain exhibit complex structure shaped by partial differentiation between *pipiens* and *molestus* ecotypes alongside extensive shared ancestry. This pattern supports the idea that *Cx. pipiens s.s.* populations in the South of the Iberian Peninsula are genetically intermediate relative to northern populations of both ecotypes. Further studies are needed to determine whether this scenario is common across Mediterranean populations. Habitat, geography, and *Wolbachia* infection further influence genetic variation, while chromosomal inversions, though not driving ecotype differentiation, may carry adaptive variants affecting vector behavior and resistance. Our findings challenge the view of hybrids as primary drivers of WNV spillover; instead, spillover risk may depend more on ecotype abundance, local ecology, and host availability. Mosquito control strategies should consider this complexity, including spatial structuring of *molestus* populations and symbiont dynamics, and winter persistence in underground breeding sites. This integrated genomic and ecological framework enhances understanding of vector population dynamics and informs targeted interventions to reduce WNV transmission risk.

## Materials and Methods

### Mosquito collection

We collected mosquitoes between May and November 2022 from 24 different localities across five provinces in Andalusia, southwestern Spain (Supplementary Table 1). Several sampling sites overlapped with areas affected by major WNV human outbreaks that occurred in 2020 and 2024. We conducted mosquito capture and morphological identification following Cebrián-Camisón *et al*. ^54^. All collections were conducted with the required permits from landowners and local authorities (Consejería de Medio Ambiente, Junta de Andalucía). After identification, mosquitoes were stored individually at -80°C until molecular analyses. Both male and female *Culex pipiens s.s* were used in this study, although females showing evidence of blood feeding were excluded.

### Molecular analyses

High-molecular-weight (HMW) genomic DNA was extracted from whole mosquitoes using the MagAttract HMW DNA Kit and DynaMagTM -2 magnet (Qiagen, Hilden, Germany) following a protocol optimized for mosquitoes ^55^. DNA concentration was measured using Qubit 3.0 Fluorometer and Qubit dsDNA HS Assay Kit (Thermofisher, Waltham, Massachusetts, USA) and purity was assessed using a Nanodrop spectrophotometer (Nanodrop Technologies, Wilmington, Delaware, USA). DNA was purified and concentrated using a 0.8-1.5X AMPure XP bead clean up (Agencourt AMPure XP Bead-Based Reagent, Beckman Coulter, Brea, California, USA). Samples with concentration ≥10ng/μl were sent to Novogene (Cambridge, UK) for shotgun library preparation and paired-end sequencing (2×150bp) on an Illumina Novaseq X platform.

### Variant calling and filtering

We aligned reads against the *Cx. pipiens* reference genome (GCA_963924435.1) ^56^, using BWA-MEM v0.7.18 ^57^. We removed PCR duplicates using Picard v.3.0.0 ^58^ and merged BAM files from multiple sequencing runs for the same sample using Samtools v1.18 ^59^. Species identity was confirmed by reconstructing mitochondrial consensus sequences for each sample using *bcftools consensu*s from BCFtools v1.19 ^60,61^. Sequences were then aligned using MAFFT v7.525 in *--auto* mode ^62^. Phylogenetic reconstruction was carried out with IQ-TREE version 2.4.0, using automatic model selection (*-m* MFP) and 1,000 ultrafast bootstrap replicates ^63^.

We called variants using *bcftools mpileup* and *bcftools call*, with a minimum alignment’s mapping quality of 20, base quality of 15, and incorporating the information on sample location. After genotyping, we filtered variants for each chromosome separately and excluded highly repetitive regions *(bcftools filter -M)* that had been previously identified using Earl Gray v5.1.1 ^64^ and a transposable element library from *Culex quinquefasciatus* ^65^. We retained biallelic SNPs with minor allele frequency ≥3%, a minimum genotype depth of 5, genotype quality of 30, and less than 20% missing genotypes. Filtered variants were concatenated across chromosomes using *bcftools concat*, resulting in 4,535,359 SNPs (“full dataset”). To minimize the effect of missing data and chromosomal inversions, we generated a reduced dataset from chromosome arms 2p and 2q. First, we retained variants from two arms from chromosome 2 using *bcftools view* (-r), and filtered them to only include biallelic SNPs with a minimum allele frequency of 5%. Then, we quantified per-individual missingness using VCFtools (*--missing-indv*). Individuals were ranked according to their proportion of missing genotypes and progressively removed, one at a time, starting with those with the highest missingness. At each step, we evaluated the number of variant sites retained when requiring complete data across all remaining individuals (i.e., no missing genotypes). This iterative procedure was used to identify an optimal balance between the number of retained individuals and the number of informative sites. Based on this balance, we retained a subset of 90 individuals. This subset retained 50,170 SNPs for chromosome arm 2p and 54,954 for arm 2q, and was used for phylogenetic reconstruction, *f_3_* analyses and several PCA analyses.

Finally, in order to analyze divergence estimates over the same subset of samples, we developed a third dataset including the three chromosomes (hereafter the “reference” dataset). This dataset retained high-quality SNPs and invariant sites for the same 90 individuals. To obtain it, we used *bcftools view* to exclude non-SNP variants (-e ‘TYPE=’indel’ | TYPE=’mnp’) and filtered by genotype depth (≥5) and quality (≥30). We also retained only the target 90 individuals (using -S), filtering missing alternative alleles (-a) and sites with uncalled genotypes (-U). Then, we removed sites with any missing data (-e ‘F_MISSING>0’) and with more than two alleles (-M 2) and minor allele frequency under 5% in the case of SNPs. This reference dataset comprised 394,717 biallelic SNPs and 35,771,515 invariant sites.

All bioinformatic analyses were performed on a combination of the Galicia Supercomputing Center (CESGA) and University of Oregon HPC clusters and on dedicated Linux servers maintained at Kern-Ralph co-laboratory from University of Oregon and Doñana’s Singular Scientific-Technical Infrastructure (ICTS-RBD). More details about the parameters used in the analyses can be found in https://github.com/socebrian/pipiens_popgen.

### Population structure analysis

Using the “full” dataset, we ran PLINK v1.9 ^66^ to prune highly linked SNPs (*--indep-pairwise 50 10 0.5*), following Aardema *et al*. ^67^, and to perform a Principal Component Analysis (PCA). To make sure that the pattern found was not caused by highly related individuals that might have been captured in the same trap we carried out a PCA accounting for relatedness using a kinship matrix. We estimated a kinship matrix using the R package SNPRelate v1.40.0 ^68^ converted the matrix to a *GenotypeData* object using the GWASTools package v1.52.0 ^69^ and ran a PCA using the function *pcair* in the GENESIS package v2.36.0 ^70^. We set the kinship threshold at 0.125 to exclude parent-offspring and full siblings from the analyses. This method is robust to relatedness as the PCA is performed using only a subset of unrelated samples. Then, PCA values for the related subset of samples that were excluded are predicted from genetic similarity.

To estimate the number of genetic clusters and individual ancestry proportions we ran admixture analyses using ADMIXTURE v1.3.0 ^71^ with *k* values ranging from 1 to 8 PCA. Admixture plots were generated in R v4.4.1 ^72^ using the ggplot2 package v3.5.1^73^.

To study if the genetic structure was related to the presence of ecotypes *pipiens* and *molestus*, we performed the PCA including 48 samples that had been assigned to either ecotype (22 *molestus* and 26 *pipiens*) from Haba *et al*.^16^ (Supplementary Table 2). We downloaded the sequences from NCBI using *prefetch* from SRA toolkit v3.2.1 and then used *fasterq-dump* to convert .*sra* files to *fastq* files. The mapping and variant calling for these samples were performed separately following the same criteria used for our *Cx. pipiens s.s* samples. Before performing variant filtering, we used *bcftools merge* to merge both datasets and performed the filtering for all samples using the same parameters and thresholds used for previous analyses. We then reran PCA and ADMIXTURE analyses. We identified ecotypes in our samples by comparing PCA grouping and admixture clusters from our samples with the ones from Haba *et al*.^16^.

### Identification of chromosomal inversions

To further explore genetic variation along the genome without pre-assignment of samples to different groups we conducted a windowed PCA using WinPCA v1.1 ^74^. We ran the PCA on windows of 100 kb. To identify regions with reduced recombination, we extracted linkage disequilibrium metrics along chromosomes excluding one variant from each pair that was closer than 10 kb using PLINK and plotted the R^2^ values. WinPCA and linkage disequilibrium metrics showed indications of possible chromosomal inversions on chromosomes 1 and 3. We determined the approximate location of inversion breakpoints by direct visualization of winPCA dynamic output plots. To obtain further evidence of those inversions, we first performed PCAs restricted to those regions. We extracted regions shorter than the estimated inversion spans in order to minimize uncertainty around breakpoint locations (chromosome 1: 88-100Mbp for 1q; chromosome 3: 4.8-17.5Mbp for 3p and 150-160Mbp for 3q). Based on the structure shown, we then assigned samples to 3 groups in each inversion, A/A, a/A, a/a (stating A/A as the most frequent arrangement), and used them as a proxy for karyotype, as we could not karyotype the inversions. Using pixy version 2.0.0.beta12 ^75,76^, we obtained F_ST_ estimates (*--stats fst*) calculated on 10 kbp windows (*--window_size*) for each chromosome separately. We specified the population assignments based on the karyotype proxies (*--populations*).

### Divergence, gene flow and phylogenetic reconstruction of *Culex pipiens* ecotypes

We calculated genetic diversity and divergence from the reference dataset of 90 samples using pixy. We specified the population assignments based on ecotype identity using the population map (*--populations*). Using the *--stats* argument, we calculated the nucleotide diversity (π) within each group, as well as the absolute (D_XY_), and relative genetic divergence (F_ST_) between groups. These statistics were calculated for 10 kbp and 5kb windows (*--window_size*), independently for each chromosome, and using only the target sites retained in the filtered VCF (*--sites_file*). In addition to these estimates from pixy, we calculated Nei’s unbiased genetic distance (D_A_) ^77^ between the populations using a custom Python script (*calculate_Da_pixy.py*, available at https://github.com/socebrian/pipiens_popgen). D_A_ estimates ensure smaller biases due to small sample sizes, while accounting for the differences in nucleotide diversity between the compared populations ^78^. All the statistics were calculated twice setting different population assignments. First, we grouped by ecotype’s ancestry of 95% or higher, leaving the rest of samples as the third population. Then, we grouped by ecotype (60%-100% ancestry), setting unassigned samples to any of them (40-60% ancestry) as the third group. We did this in order to assess the robustness of genomic patterns to the ecotype definition, given the large proportion of samples exhibiting mixed ancestry.

To establish the phylogenetic relationships between the ecotypes and unassigned individuals, we used fastreeR version 2.0.0 software ^79^ to calculate a neighbor joining tree based on the pairwise genetic distance between the 90 *Culex* individuals. We calculated two phylogenies based on the genetic information present in each arm for *Culex* chromosome 2. We restricted the analysis to chromosome 2, as this chromosome shows no evidence of large chromosomal inversions in our dataset. We used the *bcftools prune* plugin to retain only a single variant site per 1 kbp (*-n 1 -w 1000*) from the dataset of biallelic SNPs from chromosome arms 2p and 2q (N=90). By default, we pruned to retain the variant site with the highest allele frequency within the 1 kbp window (*-N ‘maxAF’*). After filtering the VCF, we used the *VCF2TREE* command in fastreeR to calculate the pairwise distance matrix for all individuals and generate the output tree in Newick format. We then plotted the tree using the *ggtree* R package v3.15.0 ^80^.

Finally, to test whether individuals with intermediate ecotype ancestry represent potential admixture between *pipiens* and *molestus*, we calculated *f_3_* statistics using *qp3Pop* from ADMIXTOOLS v7.0.1 ^81^. This analysis evaluates whether a target population shows excess shared genetic drift with two source populations, which is indicative of admixture. Prior to analysis, SNPs from chromosome 2 were pruned for linkage disequilibrium using PLINK (*--indep-pairwise 200 10 0.2*). Genotype data were converted from to EIGENSTRAT format using *convert,* from ADMIXTOOLS. Population labels were assigned grouping individuals into *pipiens*, *molestus*, and *unassigned* (40–60% ancestry). The populations file followed the standard three-column EIGENSTRAT with individual ID, sex (set to U, for unknown) and population label (*pipiens*, *molestus* or unassigned). We next specified the *f₃* test configurations listing each combination of target and source populations between *pipiens, molestus* and *unassigned*. Finally, we run *qp3Pop,* enabling estimation of f₃ for each configuration under an inbreeding-aware model (*inbreed: YES*).

### Annotation and enrichment analyses

In order to perform functional enrichment analyses, we first annotated *Cx. pipiens* reference genome (GCA_963924435.1)^82^, as the assembly does not have an associated annotation (details on Supplementary material). We then performed Gene Ontology (GO) enrichment analysis restricted to chromosomal inversions. Candidate genes per inversion were extracted using *bcftools intersect*. We limited the search to the same chromosome spans we used for the PCA analyses inside chromosomal inversions in order to minimize uncertainty around breakpoints locations. We also conducted enrichment analyses between *pipiens* and *molestus* samples (>95% ecotype ancestry), targeting 10kb and 5kb windows with the 1% higher values of F_ST_ between ecotypes, excluding chromosomal inversions. Enrichment analyses were performed following the pipeline from Lorenzo-Fernández ^83^, which uses topGO R package v2.58.0 ^84^. To correct for multiple testing, we applied the Benjamini–Hochberg procedure (BH) to control the false discovery rate (FDR). For each GO term, adjusted p-values were calculated from the original Fisher’s exact test p-values and produced filtered datasets retaining only those terms significant at α = 0.05 after correction.

### Characterization of infection by *Wolbachia*

To identify infection by *Wolbachia* we carried out competitive mapping against a combined *Cx. pipiens* and *Wolbachia pipientis* (GCA_000073005.1) reference using the same mapping and read curation protocol as for the previous alignments with *Cx. Pipiens s.s*. To classify the samples as *w*Pip^+^ or *w*Pip^-^, we extracted paired reads with at least 60% of their length aligned against the *Wolbachia* genome and used a custom Python script (Alice Namias and Myléne Weill, personal communication) to estimate the percentage of *Wolbachia* genome covered. We set a conservative threshold of 1% of genome coverage to consider a *Cx. pipiens* as *w*Pip^+^. To identify the *w*Pip group (*w*Pip-I to *w*Pip-V) infecting the mosquitoes we used the *pk1* marker, a fragment of approximately 1300bp long of the ANK domain gene (*pk1*) that reliably discriminates among the five main *wPip* groups ^29,37,85^. To do this, we obtained *Wolbachia* consensus sequences using *bcftools consensu*s and kept sequences with less than 10% missing sites. To identify *p*k*1* marker on our samples, we generated a database from consensus sequences using *makeblastdb* from BLAST v.2.17.0 ^86^ and then performed BLAST using *blastn* with a set e-value of 1^- 20^ and a curated set of *pk1* reference sequences representing the five *w*Pip groups (Supplementary Table 3). Because the *pk1* sequence was identical for all our *w*Pip^+^ samples we did not perform downstream phylogenetic reconstruction.

### Isolation-by-distance and dbRDA analyses

We analyzed the relationship between genetic variance and several environmental and spatial factors in addition to ecotype differentiation. To do so we performed distance- based redundancy analyses (dbRDA) and quantified the genetic variance explained by ecotype identity, infection by *Wolbachia*, geographic location and habitat type (urban vs. natural). We calculated the pairwise Euclidean genetic distances (*--distance ibs square*) using PLINK. We used this IBD matrix as the response variable and the ecotype, habitat type, geographic multivariate matrix and *Wolbachia* infection status as the explanatory variables. As a proxy of ecotype, we included in the models the proportion of ancestry from Q1 from the ADMIXTURE analysis. To define the “habitat” variable for each location, we delimited a 2 km radius buffer around each trapping location using QGIS v3.34.11 ^87^. If an urban area fell inside this radius, the locality was assigned as urbanized. To account for spatial patterns, we developed a geographic multivariate matrix. For this, we first used QGIS to calculate the straight-line distance between all pairs of localities. Then we used Principal Coordinates of Neighbor Matrices (PCNM), obtained with *pcnm()* from *vegan* R package v2.6.10 ^88^, to model spatial autocorrelation across multiple scales. The five PCNM vectors obtained were combined into a single multivariate predictor matrix to evaluate spatial structure as a whole, reducing model complexity and avoiding multicollinearity. Finally, infection by *Wolbachia* was expressed as a discrete variable indicating if the mosquito was or was not infected. To ensure that PCNMs were not confounded with ecological or genetic variables (pQ1, habitat, *Wolbachia* infection), we tested for collinearity using Variance Inflation Factors (VIF; *vif.cca* function from *vegan*) and one-way ANOVAs. No significant associations were detected (all p>0.05) and all VIF values were < 2, suggesting very low collinearity among variables. Thus, we retained all predictors to perform dbRDA using the *capscale()* function in *vegan*.

To test whether the effect of Euclidean distance over genetic divergence changed between ecotypes, we performed Mantel tests with *vegan* for both ecotypes separately, using Pearson correlations and 9,999 permutations. Finally, we performed a beta generalized linear mixed-effects model (GLMM) to model the effect of habitat (natural/urban) over individual ancestry proportions. Model was fit using glmmTMB R package ^89^ with a logit link function. Location was included as a random effect.

## Acknowledgments

We are grateful to Cristina Pérez and Begoña Adrados for their help in the molecular analyses, and to Alvaro Solís for his help during mosquito collection and morphological identification. We also want to thank Lindy McBride and Yuki Haba for their input on our analyses and Alice Namias and Myléne Weill for developing the approach and script used to assess *wPip* infection from whole-genome sequence data. This work was supported by Horizon Europe under the Biodiversity, Circular Economy and Environment (REA.B.3), co-funded by the Swiss State Secretariat for Education, Research and Innovation (SERI) under contract numbers 22.00173 and 24.00054, by the UK Research and Innovation (UKRI) under the Department for Business, Energy and Industrial Strategy’s Horizon Europe Guarantee Scheme and PID2021-123761OB-I00/AEI/10.13039/501100011033/FEDER,UE. The project and MJRL were supported by the Spanish State Research Agency (project PID2020-118921RJ-100/AEI/10.13039/501100011033). SC was supported by the Spanish Ministry of Science, Innovation and Universities (FPU program, FPU20/03477) and conducted part of the research that led to this publication with the support of a US-Spain Fulbright grant co-funded by Fulbright Spain and Junta de Andalucía (grant number: 2024-84). A.D.K was supported in part by NIH award R35GM148253.

## Data Availability

The raw sequencing data generated in this study have been deposited in the European Nucleotide Archive (ENA) under accession number PRJEB108586. Sample metadata are available through COPO. The genome annotation and mosquito capture protocols are available in Zenodo [DOIs/links to be added upon acceptance]. Custom code and analysis pipelines used in this study are publicly available at https://github.com/socebrian/pipiens_popgen

## Authors contributions

M.J.R.L. and J.F. conceived and supervised the study. M.J.R.L. and S.C.C. optimized DNA extraction protocols. S.C.C. performed field mosquito collections and DNA extractions. S.C.C and A.G.R.C performed genomic and statistical analyses. A.B. developed the genome annotation for *Culex pipiens* genome assembly. A.D.K., P.L.R., A.G.R.C., S.T.S. provided input on analytical approaches and contributed to the interpretation of the results. J.F. provided expertise on species-specific biology and ecology of *Culex pipiens*. S.C.C. and M.J.R.L. were responsible for data curation and availability. A.D.K and P.L.R. provided computing resources. M.J.R.L. and J.F. secured funding for the study. S.C.C and M.J.R.L wrote the first draft of the manuscript and all the authors contributed to the revision and editing of the manuscript.

## References

1. Weissenböck, H., Hubálek, Z., Bakonyi, T. & Nowotny, N. Zoonotic mosquito-borne flaviviruses: Worldwide presence of agents with proven pathogenicity and potential candidates of future emerging diseases. Vet. Microbiol. 140, 271–280 (2010).

2. Camp, J. V. & Nowotny, N. The knowns and unknowns of West Nile virus in Europe: what did we learn from the 2018 outbreak? Expert Rev. Anti Infect. Ther. 18, 145–154 (2020).

3. ECDC. Monthly updates: Seasonal surveillance in humans and animals in 2025 for West Nile virus. https://www.ecdc.europa.eu/en/infectious-disease-topics/west-nile-virus-infection/surveillance-and-disease-data/monthly-updates (2025).

4. Taheri, S., González, M. A., Ruiz-López, M. J., Soriguer, R. & Figuerola, J. Patterns of West Nile virus vector co-occurrence and spatial overlap with human cases across Europe. One Health 20, 101041 (2025).

5. Vogels, C. B., Göertz, G. P., Pijlman, G. P. & Koenraadt, C. J. Vector competence of European mosquitoes for West Nile virus. Emerg. Microbes Infect. 6, 1–13 (2017).

6. Giesen, C. et al. A systematic review of environmental factors related to WNV circulation in European and Mediterranean countries. One Health 16, 100478 (2023).

7. Petersen, L. R., Brault, A. C. & Nasci, R. S. West Nile virus: Review of the Literature. JAMA 310, 308–315 (2013).

8. Vinogradova, E. B. Culex Pipiens Pipiens Mosquitoes: Taxonomy, Distribution, Ecology, Physiology, Genetics, Applied Importance and Control. (Pensoft, 2000).

9. Farajollahi, A., Fonseca, D. M., Kramer, L. D. & Marm Kilpatrick, A. “Bird biting” mosquitoes and human disease: A review of the role of *Culex pipiens* complex mosquitoes in epidemiology. Infect. Genet. Evol. 11, 1577–1585 (2011).

10. Haba, Y. & McBride, L. Origin and status of Culex pipiens mosquito ecotypes. Curr. Biol. 32, R237–R246 (2022).

11. Magallanes, S., Ruiz-López, M. J. & Figuerola, J. Winter ecology of the main mosquito vectors of West Nile virus in a human outbreak hotspot in southern Europe. Sci. Total Environ. 1008, 181001 (2025).

12. Chevillon, C. et al. Migration/selection balance and ecotypic differentiation in the mosquito Culex pipiens. Mol. Ecol. 7, 197–208 (1998).

13. Gomes, B. et al. Asymmetric introgression between sympatric molestus and pipiens forms of Culex pipiens (Diptera: Culicidae) in the Comporta region, Portugal. BMC Evol. Biol. 9, 262 (2009).

14. Huang, S. et al. Genetic Variation Associated with Mammalian Feeding in Culex pipiens from a West Nile virus Epidemic Region in Chicago, Illinois. Vector-Borne Zoonotic Dis. 9, 637–642 (2009).

15. Kilpatrick, A. M. et al. Genetic Influences on Mosquito Feeding Behavior and the Emergence of Zoonotic Pathogens. Am. J. Trop. Med. Hyg. 77, 667–671 (2007).

16. Haba, Y. et al. Ancient origin of an urban underground mosquito. Science 390, eady4515 (2025).

17. Dobzhansky, T. Genetics and the Origin of Species. (Columbia University Press, New York, 1937).

18. Hoffmann, A. A. & Rieseberg, L. H. Revisiting the Impact of Inversions in Evolution: From Population Genetic Markers to Drivers of Adaptive Shifts and Speciation? Annu. Rev. Ecol. Evol. Syst. 39, 21–42 (2008).

19. Bryan, J., Petrarca, V., Di Deco, M. & Coluzzi, M. Adult behaviour of members of the Anopheles gambiae complex in the Gambia with special reference to An. melas and its chromosomal variants. Parassitologia 29, (1987).

20. Riehle, M. M. et al. The Anopheles gambiae 2La chromosome inversion is associated with susceptibility to Plasmodium falciparum in Africa. eLife 6, e25813 (2017).

21. Unger, M. F., Sharakhova, M. V., Harshbarger, A. J., Glass, P. & Collins, F. H. A standard cytogenetic map of Culex quinquefasciatus polytene chromosomes in application for fine-scale physical mapping. Parasit. Vectors 8, 307 (2015).

22. Minwuyelet, A. et al. Symbiotic Wolbachia in mosquitoes and its role in reducing the transmission of mosquito-borne diseases: updates and prospects. Front. Microbiol. 14, (2023).

23. Werren, J. H., Baldo, L. & Clark, M. E. Wolbachia: master manipulators of invertebrate biology. Nat. Rev. Microbiol. 6, 741–751 (2008).

24. Garrigós, M., Garrido, M., Panisse, G., Veiga, J. & Martínez-de la Puente, J. Interactions between West Nile virus and the Microbiota of Culex pipiens Vectors: A Literature Review. Pathogens 12, 1287 (2023).

25. Brucker, R. M. & Bordenstein, S. R. Speciation by symbiosis. Trends Ecol. Evol. 27, 443–451 (2012).

26. Vavre, F., Fleury, F., Varaldi, J., Fouillet, P. & Boulétreau, M. Infection polymorphism and cytoplasmic incompatibility in Hymenoptera-Wolbachia associations. Heredity 88, 361–365 (2002).

27. Bergman, A. & Hesson, J. C. Wolbachia prevalence in the vector species Culex pipiens and Culex torrentium in a Sindbis virus-endemic region of Sweden. Parasit. Vectors 14, 428 (2021).

28. Yang, Y., et al. Prevalence and molecular characterization of Wolbachia in field-collected Aedes albopictus, Anopheles sinensis, Armigeres subalbatus, Culex pipiens and Cx. tritaeniorhynchus in China. PLoS Negl. Trop. Dis. 15, e0009911 (2021).

29. Dumas, E. et al. Population structure of Wolbachia and cytoplasmic introgression in a complex of mosquito species. BMC Evol. Biol. 13, 181 (2013).

30. Lilja, T., Lindström, A., Hernández-Triana, L. M., Di Luca, M. & Lwande, O. W. European Culex pipiens Populations Carry Different Strains of Wolbachia pipientis. Insects 15, 639 (2024).

31. McCoy, K. D. The population genetic structure of vectors and our understanding of disease epidemiology. Parasite 15, 444–448 (2008).

32. Figuerola, J., Soriguer, R., Rojo, G., Tejedor, C. G. & Jimenez-Clavero, M. A. Seroconversion in Wild Birds and Local Circulation of West Nile virus, Spain - Volume 13, Number 12—December 2007 - Emerging Infectious Diseases journal - CDC. Emerg. Infect. Dis. 13, 1915–1917 (2007).

33. Ministerio de Sanidad. Meningoencefalitis Por Virus Del Nilo Occidental En España. Resumen de La Temporada 2024. https://www.sanidad.gob.es/areas/alertasEmergenciasSanitarias/preparacionRespuesta/docs/20250131_ERR_Nilo_Occidental.pdf (2025).

34. Rodríguez-Alarcón, L. G. S. M. et al. Unprecedented increase of West Nile virus neuroinvasive disease, Spain, summer 2020. Eurosurveillance 26, 2002010 (2021).

35. Figuerola, J. et al. A One Health view of the West Nile virus outbreak in Andalusia (Spain) in 2020. Emerg. Microbes Infect. 11, 2570–2578 (2022).

36. Martínez-de la Puente, J., et al. Culex pipiens forms and urbanization: effects on blood feeding sources and transmission of avian Plasmodium. Malar. J. 15, 589 (2016).

37. Atyame, C. M., Delsuc, F., Pasteur, N., Weill, M. & Duron, O. Diversification of Wolbachia Endosymbiont in the Culex pipiens Mosquito. Mol. Biol. Evol. 28, 2761–2772 (2011).

38. Ding, H., Yeo, H. & Puniamoorthy, N. Wolbachia infection in wild mosquitoes (Diptera: Culicidae): implications for transmission modes and host-endosymbiont associations in Singapore. Parasit. Vectors 13, 612 (2020).

39. Marques, D. A., Meier, J. I. & Seehausen, O. A Combinatorial View on Speciation and Adaptive Radiation. Trends Ecol. Evol. 34, 531–544 (2019).

40. Johannesson, K., Malmqvist, G., Leder, E. & Stankowski, S. Genomic insights into the origin of ecotypes. Trends Ecol. Evol. S0169534725003453 (2025) doi:10.1016/j.tree.2025.11.011.

41. Coutinho-Abreu, I. V., Riffell, J. A. & Akbari, O. S. Human attractive cues and mosquito host-seeking behavior. Trends Parasitol. 38, 246–264 (2022).

42. Foster, W. A. Mosquito Sugar Feeding and Reproductive Energetics. Annu. Rev. Entomol. 40, 443–474 (1995).

43. Suh, E., Choe, D.-H., Saveer, A. M. & Zwiebel, L. J. Suboptimal Larval Habitats Modulate Oviposition of the Malaria Vector Mosquito Anopheles coluzzii. PLOS ONE 11, e0149800 (2016).

44. Gadji, M. et al. Genomic Drivers of Pyrethroid Resistance Escalation in the Malaria Vector Anopheles funestus Across Africa. Mol. Biol. Evol. 42, msaf251 (2025).

45. Kefi, M. et al. Transcriptomic analysis of resistance and short-term induction response to pyrethroids, in Anopheles coluzzii legs. BMC Genomics 22, 891 (2021).

46. Pignatelli, P. et al. The Anopheles gambiae ATP-binding cassette transporter family: phylogenetic analysis and tissue localization provide clues on function and role in insecticide resistance. Insect Mol. Biol. 27, 110–122 (2018).

47. Yang, T. & Liu, N. Genome Analysis of Cytochrome P450s and Their Expression Profiles in Insecticide Resistant Mosquitoes, Culex quinquefasciatus. PLOS ONE 6, e29418 (2011).

48. Huang, Y. et al. Culex pipiens pallens cuticular protein CPLCG5 participates in pyrethroid resistance by forming a rigid matrix. Parasit. Vectors 11, 6 (2018).

49. Kwiatkowska, R. M. et al. Dissecting the mechanisms responsible for the multiple insecticide resistance phenotype in *Anopheles gambiae* s.s., M form, from Vallée du Kou, Burkina Faso. Gene 519, 98–106 (2013).

50. Fritz, M. L., Walker, E. D., Miller, J. R., Severson, D. W. & Dworkin, I. Divergent host preferences of above- and below-ground Culex pipiens mosquitoes and their hybrid offspring. Med. Vet. Entomol. 29, 115–123 (2015).

51. Osório, H. C., Zé-Zé, L., Amaro, F., Nunes, A. & Alves, M. J. Sympatric occurrence of Culex pipiens (Diptera, Culicidae) biotypes pipiens, molestus and their hybrids in Portugal, Western Europe: feeding patterns and habitat determinants. Med. Vet. Entomol. 28, 103–109 (2014).

52. Gomes, B. et al. Feeding patterns of molestus and pipiens forms of Culex pipiens (Diptera: Culicidae) in a region of high hybridization. Parasit. Vectors 6, 93 (2013).

53. Rizzoli, A. et al. Understanding West Nile virus ecology in Europe: Culex pipiens host feeding preference in a hotspot of virus emergence. Parasit. Vectors 8, 213 (2015).

54. Cebrián-Camisón, S., Ruiz-Lopez, M., Magallanes, S. & Figuerola, J. Mosquito collection and handling protocol to obtain High-Molecular Weight DNA. 10.5281/zenodo.20072157 (2026).

55. Teltscher, F., Johnson, H. & Lawniczak, M. Manual extraction of High Molecular Weight DNA from single mosquitoes using the Qiagen MagAttract HMW DNA kit. https://www.protocols.io/view/manual-extraction-of-high-molecular-weight-dna-fro-cra7v2hn (2023).

56. Hesson, J. C. et al. A chromosomal reference genome sequence for the northern house mosquito, Culex pipiens form pipiens, Linnaeus, 1758. Wellcome Open Res. 10, 107 (2025).

57. Li, H. & Durbin, R. Fast and accurate short read alignment with Burrows–Wheeler transform. Bioinformatics 25, 1754–1760 (2009).

58. Broad Institute. Picard Toolkit. https://github.com/broadinstitute/picard (2019).

59. Li, H. et al. The Sequence Alignment/Map format and SAMtools. Bioinformatics 25, 2078–2079 (2009).

60. Danecek, P. et al. Twelve years of SAMtools and BCFtools. GigaScience 10, giab008 (2021).

61. Li, H. A statistical framework for SNP calling, mutation discovery, association mapping and population genetical parameter estimation from sequencing data. Bioinformatics 27, 2987–2993 (2012).

62. Katoh, K. & Standley, D. M. MAFFT Multiple Sequence Alignment Software Version 7: Improvements in Performance and Usability. Mol. Biol. Evol. 30, 772–780 (2013).

63. Minh, B. Q. et al. IQ-TREE 2: New Models and Efficient Methods for Phylogenetic Inference in the Genomic Era. Mol. Biol. Evol. 37, 1530–1534 (2020).

64. Baril, T., Galbraith, J. & Hayward, A. Earl Grey: A Fully Automated User-Friendly Transposable Element Annotation and Analysis Pipeline. Mol. Biol. Evol. 41, msae068 (2024).

65. Ryazansky, S. S. et al. The chromosome-scale genome assembly for the West Nile vector Culex quinquefasciatus uncovers patterns of genome evolution in mosquitoes. BMC Biol. 22, 16 (2024).

66. Purcell, S. et al. PLINK: A Tool Set for Whole-Genome Association and Population-Based Linkage Analyses. Am. J. Hum. Genet. 81, 559–575 (2007).

67. Aardema, M. L., vonHoldt, B. M., Fritz, M. L. & Davis, S. R. Global evaluation of taxonomic relationships and admixture within the Culex pipiens complex of mosquitoes. Parasit. Vectors 13, 8 (2020).

68. Zheng, X. et al. A high-performance computing toolset for relatedness and principal component analysis of SNP data. Bioinformatics 28, 3326–3328 (2012).

69. Gogarten, S. M. et al. GWASTools: an R/Bioconductor package for quality control and analysis of genome-wide association studies. Bioinformatics 28, 3329–3331 (2012).

70. Gogarten, S. M. et al. Genetic association testing using the GENESIS R/Bioconductor package. Bioinformatics 35, 5346–5348 (2019).

71. Alexander, D. H., Novembre, J. & Lange, K. Fast model-based estimation of ancestry in unrelated individuals. Genome Res. 19, 1655–1664 (2009).

72. R Core Team. R: A Language and Environment for Statistical Computing. https://www.R-project.org/ (2025).

73. Wickham, H. Ggplot2. (Springer International Publishing, Cham, 2016). doi:10.1007/978-3-319-24277-4.

74. Blumer, L. M., Good, J. M. & Durbin, R. WinPCA: A package for windowed principal component analysis. Preprint at 10.48550/arXiv.2501.11982 (2025).

75. Bailey, N., Stevison, L. & Samuk, K. Correcting for Bias in Estimates of θw and Tajima’s D From Missing Data in Next-Generation Sequencing. Mol. Ecol. Resour. 25, e14104 (2025).

76. Korunes, K. L. & Samuk, K. pixy: Unbiased estimation of nucleotide diversity and divergence in the presence of missing data. Mol. Ecol. Resour. 21, 1359–1368 (2021).

77. Nei, M. ESTIMATION OF AVERAGE HETEROZYGOSITY AND GENETIC DISTANCE FROM A SMALL NUMBER OF INDIVIDUALS. Genetics 89, 583–590 (1978).

78. Hahn, M. W. Molecular Population Genetics. (Oxford University Press, New York, NY, 2019).

79. Gkanogiannis, A., Gazut, S., Salanoubat, M., Kanj, S. & Brüls, T. A scalable assembly-free variable selection algorithm for biomarker discovery from metagenomes. BMC Bioinformatics 17, 311 (2016).

80. Xu, S., et al. Ggtree: A serialized data object for visualization of a phylogenetic tree and annotation data. iMeta 1, e56 (2022).

81. Patterson, N. et al. Ancient Admixture in Human History. Genetics 192, 1065–1093 (2012).

82. Bombarely, A., Cebrián-Camisón, S. & Ruiz-Lopez. Genome annotation of Culex pipiens assembly idCulPipi1.1 (GCA_963924435.1). Zenodo 10.5281/zenodo.19707072 (2026).

83. Lorenzo-Fernández, L. GO Enrichment Analysis Pipeline. https://github.com/lorenalorenzo/Functional_enrichment/tree/main?tab=readme-ov-file#go-enrichment-analysis-pipeline (2025).

84. Alexa, A. & Rahnenfuhrer, J. TopGO: Enrichment Analysis for Gene Ontology. (2025).

85. da Moura, A. J. F. et al. Screening of natural Wolbachia infection in mosquitoes (Diptera: Culicidae) from the Cape Verde islands. Parasit. Vectors 16, 142 (2023).

86. Camacho, C. et al. BLAST+: architecture and applications. BMC Bioinformatics 10, 421 (2009).

87. QGIS.org. QGIS Geographic Information System. (2025).

88. Oksanen, J., et al. vegan: Community Ecology Package. 2.7–1 10.32614/CRAN.package.vegan (2025).

89. Brooks, M. E. et al. glmmTMB Balances Speed and Flexibility Among Packages for Zero-inflated Generalized Linear Mixed Modeling. R J. 9, 378–400 (2017).

