## Supplementary Material for "Ecotypes, *Wolbachia,* and urbanization shape *Culex pipiens* population structure in a West Nile virus hotspot"

**Supplementary Methods**

**Genome annotation**

The assembly for *Culex pipiens pipiens* (accession number GCA_963924435.1) was downloaded from NCBI genome database. The seqIDs were replaced by new seqIDs containing the number of the chromosomes using custom Perl one-line (perl -ne) commands (https://github.com/socebrian/pipiens_popgen/blob/main/6.Annotation_and_enrichment.md). A mapping file was created with old and new sequence IDs. The assembly was evaluated with QUAST v5.2.0 (Gurevich et al., 2013) and BUSCO v5.6.1 (diptera_odb10 dataset) (Simão et al., 2015). Repetitive elements were annotated with the EarlGrey pipeline v5.1.1 (Baril et al., 2024) with the default parameters, rDNA elements were annotated with Barrnap v0.9. (Seemann, 2025) and tDNA elements were annotated with tRNAscan-SE v2.0.12 (Chan et al., 2021).

Evidence-based gene structural annotation was performed with transcriptomic and protein sequence data. In brief, 36 transcriptomic datasets (RNA-Seq) were downloaded from EBI database (study ERX10251346). They were processed with Fastq-mcf v1.04.676 (q ≥30, length ≥50 bp) (Aronesty, 2013) and mapped to the reference assembly with STAR v2.7.11b (Dobin et al., 2013), with default parameters compatible with StringTie . We generated transcript models with StringTie v2.2.3 (Pertea et al., 2015) and identified CDS with TransDecoder v5.7.1 (Haas, 2025). We downloaded genome assemblies and annotations for 15 species (GenBank assembly IDs between parenthesis) from NCBI: *Aedes aegypti* (GCF_002204515.2), *Aedes albopictus* (GCF_035046485.1), *Anopheles funestus* (GCF_943734845.2), *Anopheles gambiae* (GCF_943734735.2), *Anopheles stephensi* (GCF_013141755.1), *Armigeres subalbatus* (GCF_024139115.2), *Culex pipiens subsps. pallens* (GCF_016801865.2), *Culex quinquefasciatus* (GCF_015732765.1), *Malaya genurostris* (GCF_037179485.1), *Ochlerotatus camptorhynchus* (GCF_037179485.1), *Sabethes cyaneus* (GCF_943734655.1), *Topomyia yanbarensis* (GCF_030247195.1), *Toxorhynchites rutilus subsp. septentrionalis* (GCF_029784135.1*)*, *Uranotaenia lowii* (GCF_029784155.1) and *Wyeomyia smithii* (GCF_029784165.1). We then used these data and GeMoMa v1.9 (Keilwagen et al., 2016) to perform two functional annotations, one using *Culex genomes* and another using all Culicidae data.

We used BRAKER3 (Gabriel et al., 2024) and EviAnn v2.0.2 (Zimin et al., 2025) pipelines for genome annotation. BRAKER was run with RNA-Seq only (Braker1), proteins only (Braker2), and both datasets (Braker3). EviAnn was run with both RNA-Seq and closely related proteins datasets. We selected EviAnn annotation as the best performing annotation after evaluating them with GAQET2 (https://github.com/victorgcb1987/GAQET2, in prep). Functional annotation was performed by sequence homology of the protein sequences identified with EviAnn using

Diamond v2.0.14.152 against SwissProt and TrEMBL datasets (Uniprot, Feb 2025), and curated with AHRD v3.11 (https://github.com/groupschoof/AHRD).

### Supplementary Figures

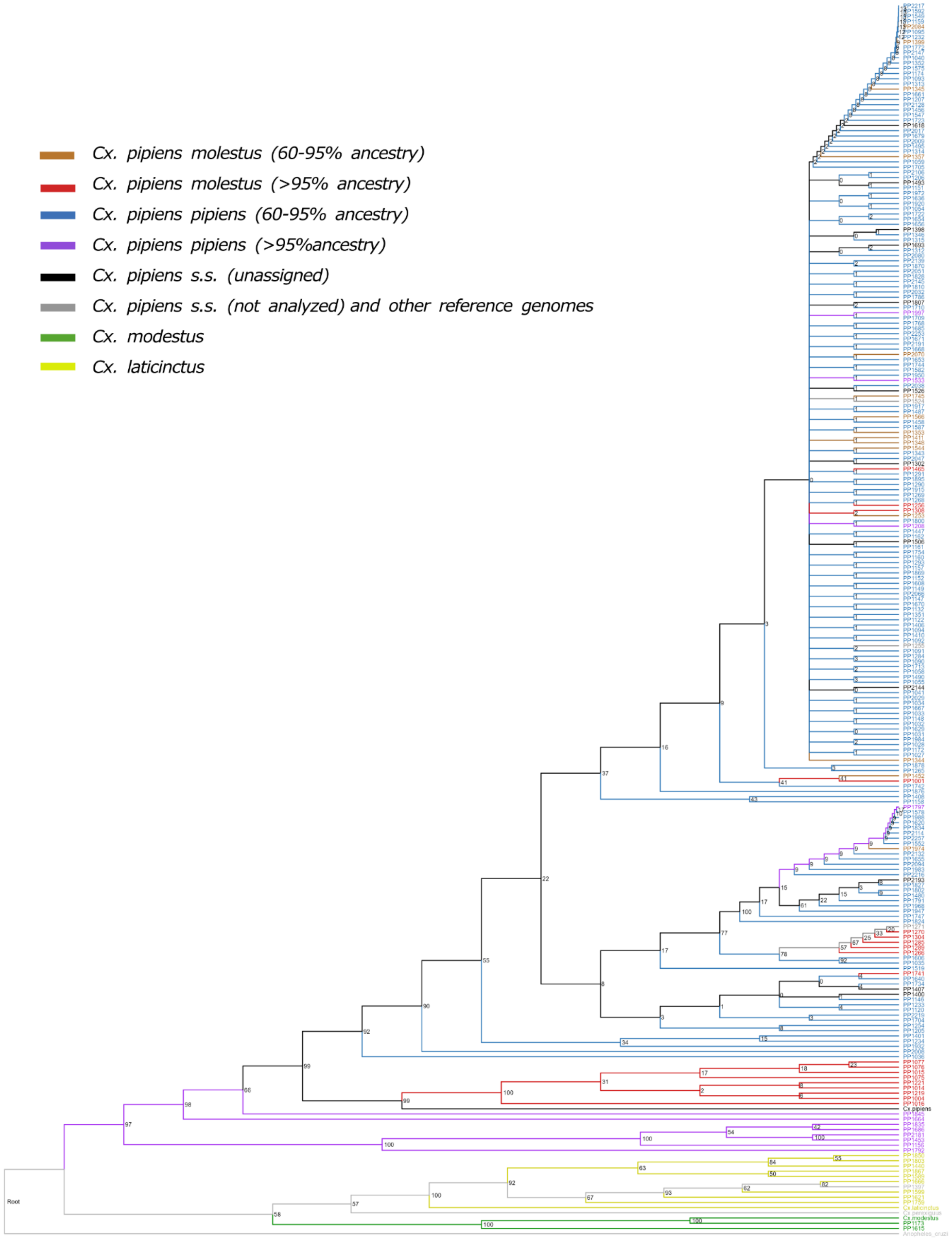

**Supplementary Figure 1**: Phylogenetic reconstruction based on mitochondrial DNA of 231 *Culex* mosquitoes from southern Spain and *Culex spp.* reference genomes. *Cx. p. pipiens* are colored in blue (60-95% *Cx. p. pipiens* ancestry) and purple (≥95% *Cx. p. pipiens* ancestry). *Cx. p. molestus* are colored in orange (60-95% *Cx. p. molestus* ancestry) and red (≥95% *Cx. p. molestus* ancestry). *Cx. pipiens s.s.* unassigned to ecotype (40-60% ecotype ancestries) and *Cx. p. pipiens* reference genome are shown in black. Ecotype assignments are based on nuclear DNA. Samples belonging to other *Culex* species, their reference genomes, and *Cx. pipiens s.s.* samples not included in further analyses because of high missing data are shown in grey.

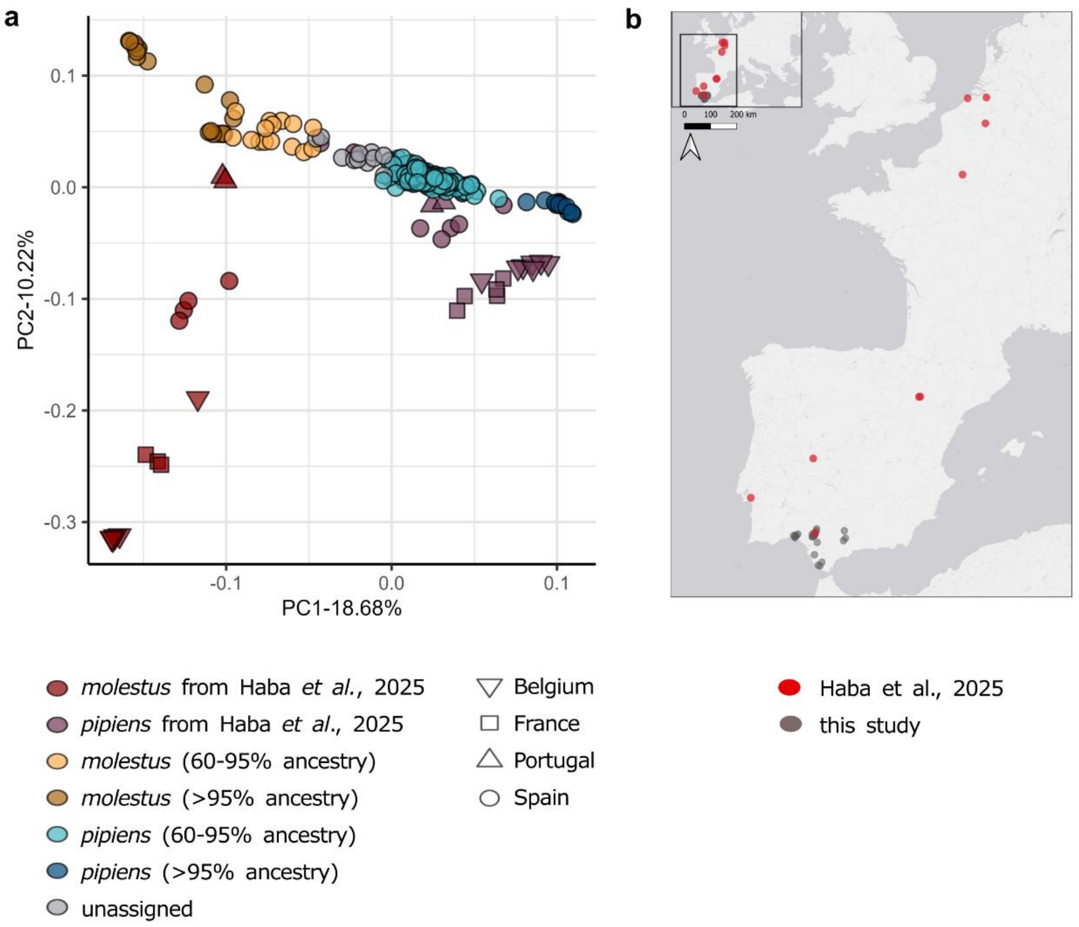


**Supplementary Figure 2**: Principal Components Analysis (PCA) including *Culex pipiens* from the present study (N=217) and from Haba *et al*. (2025, N=48). a) PCA results showing ecotype assignment. b) Map of trapping locations of samples included in the analyses.

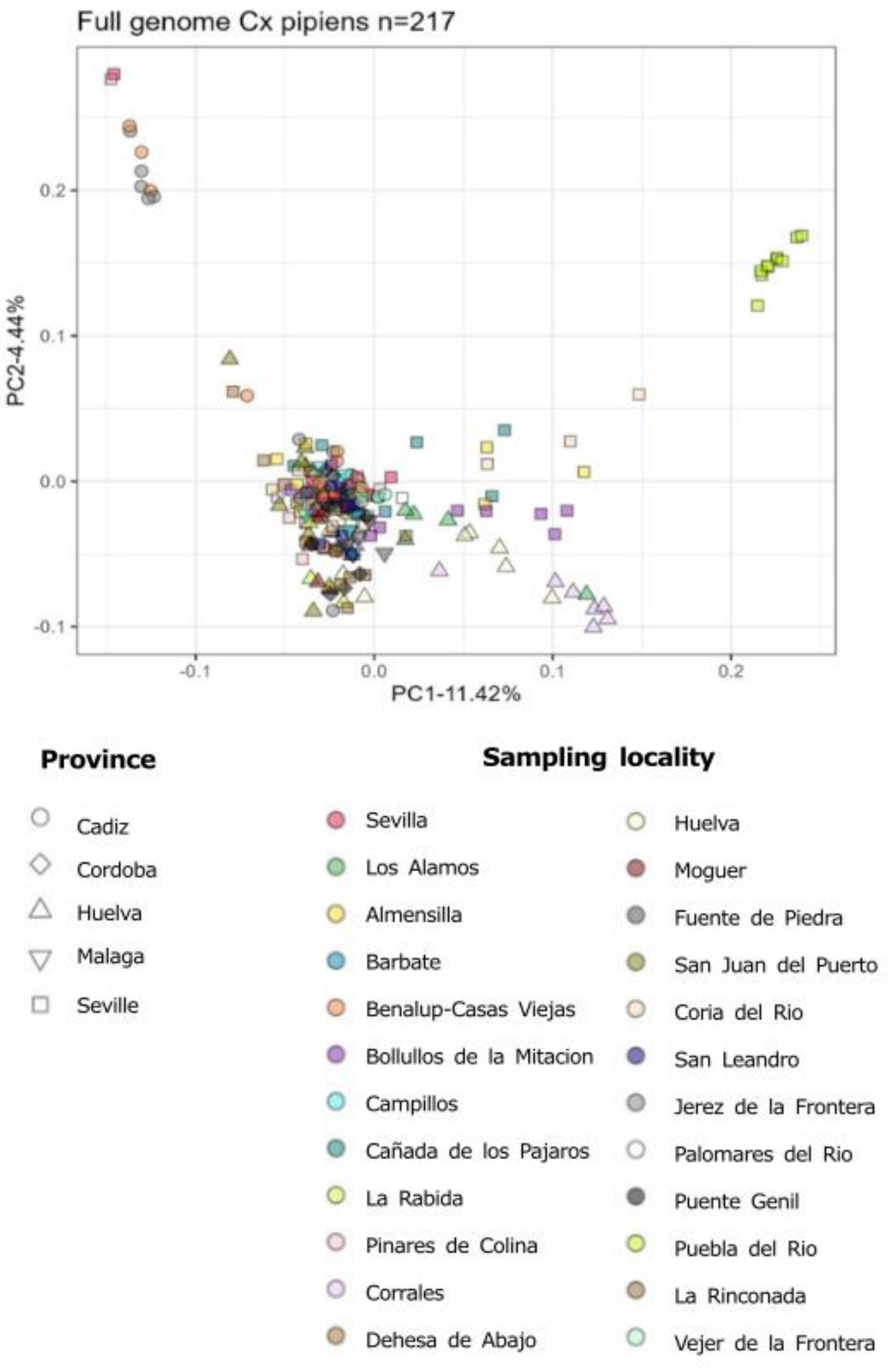


**Supplementary Figure 3**: Principal Components Analysis (PCA) of 217 *Culex pipiens* accounting for relatedness between samples (kinship threshold = 0.125).

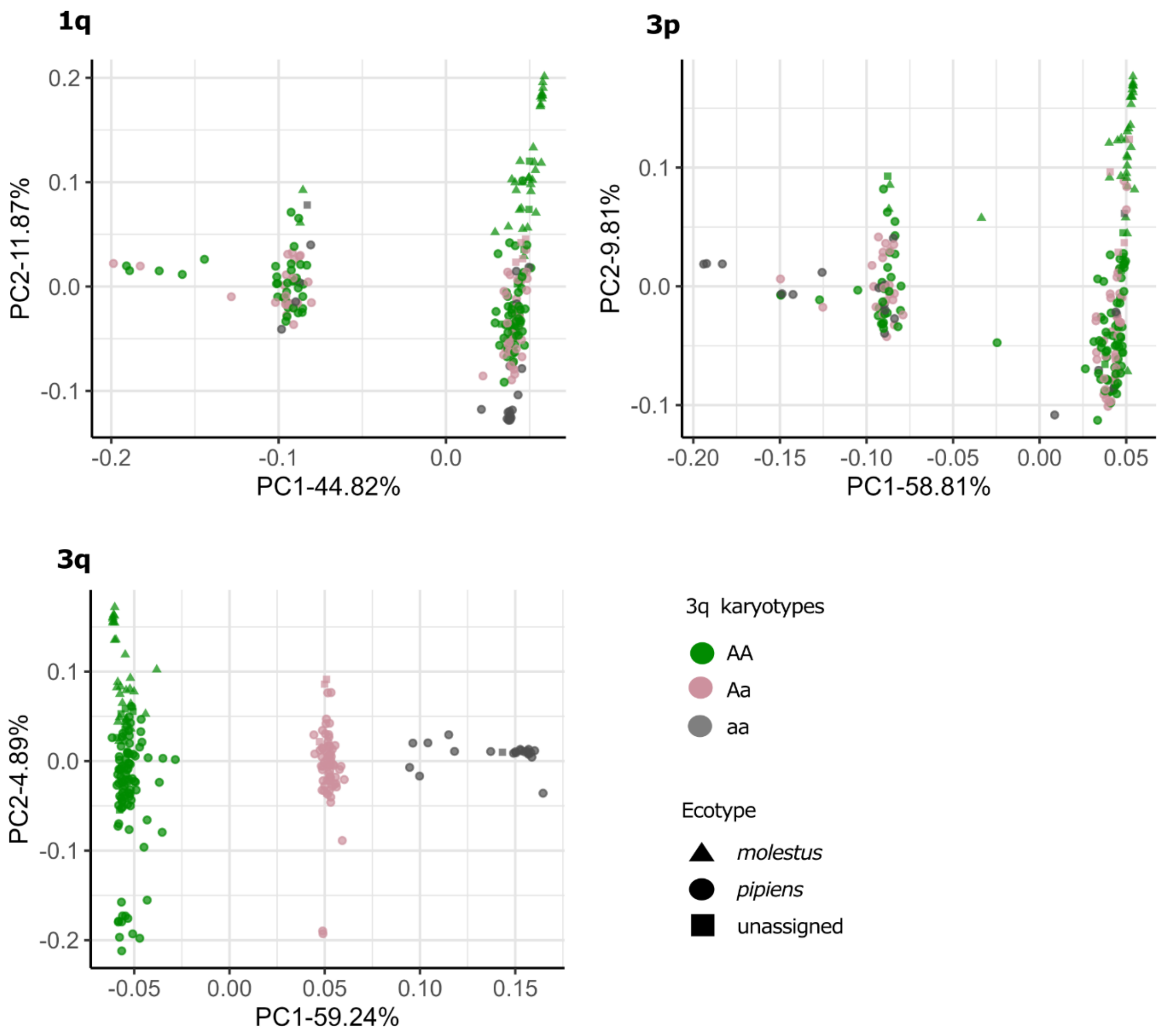


**Supplementary Figure 4:** Principal Components Analysis (PCA) of 217 *Culex pipiens* restricted to regions inside putative chromosomal inversions 1q (88-100Mbp, chromosome 1), 3p (4.8-17.5Mbp, chromosome 3) and 3q (150-160Mbp, chromosome 3). Samples are colored based on the karyotype they show in inversion 3q.

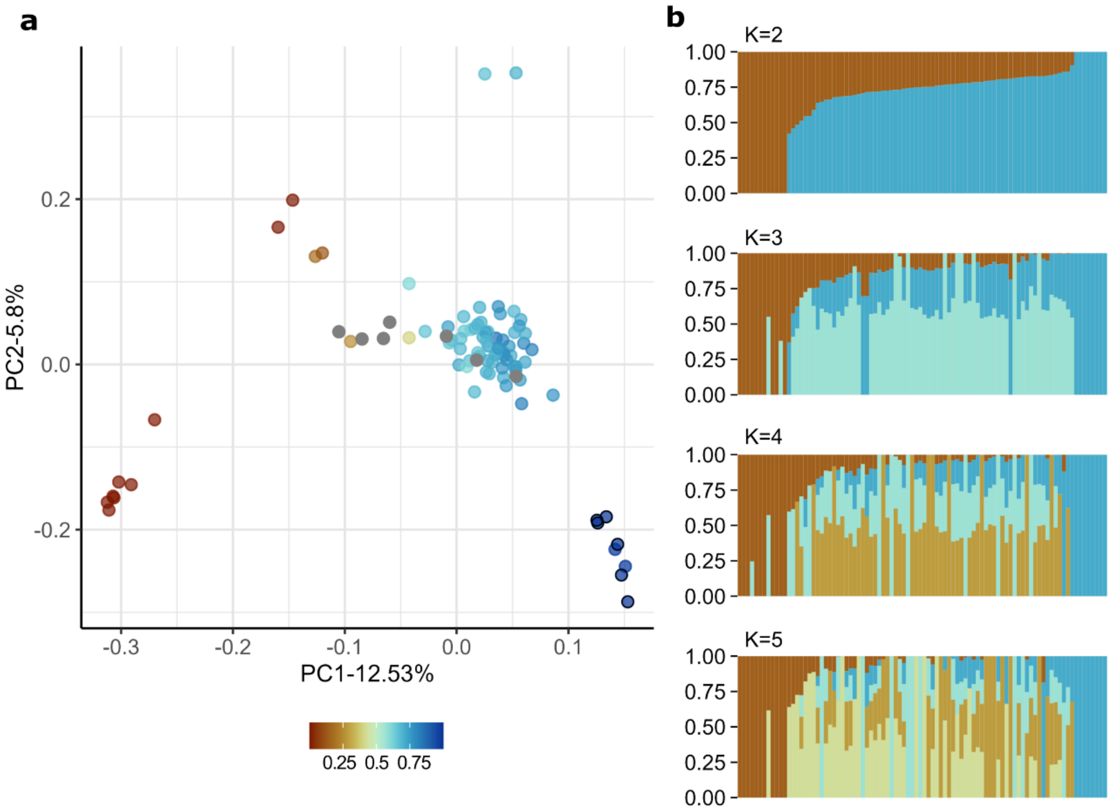


**Supplementary Figure 5:** Population structure analyses of *Culex pipiens* controlling for the confounding effect of chromosomal inversions and missing genotypes. a) Principal Components Analysis (PCA) of 90 *Cx. pipiens s.s.* Color gradient represents estimated proportion of *Cx. p. pipiens* ancestry (p(Q2) in ADMIXTURE analyses with K=2): samples with higher *Cx. p. pipiens* ancestry are shown in blue, whereas those with higher *Cx. p. molestus* ancestry are shown in orange. Grey dots represent samples that could not be confidently assigned to any of the ecotypes. Samples uninfected by Wolbachia are represented with a black outline. b) ADMIXTURE results for K=2 to K=5, illustrating individual ancestry proportions.

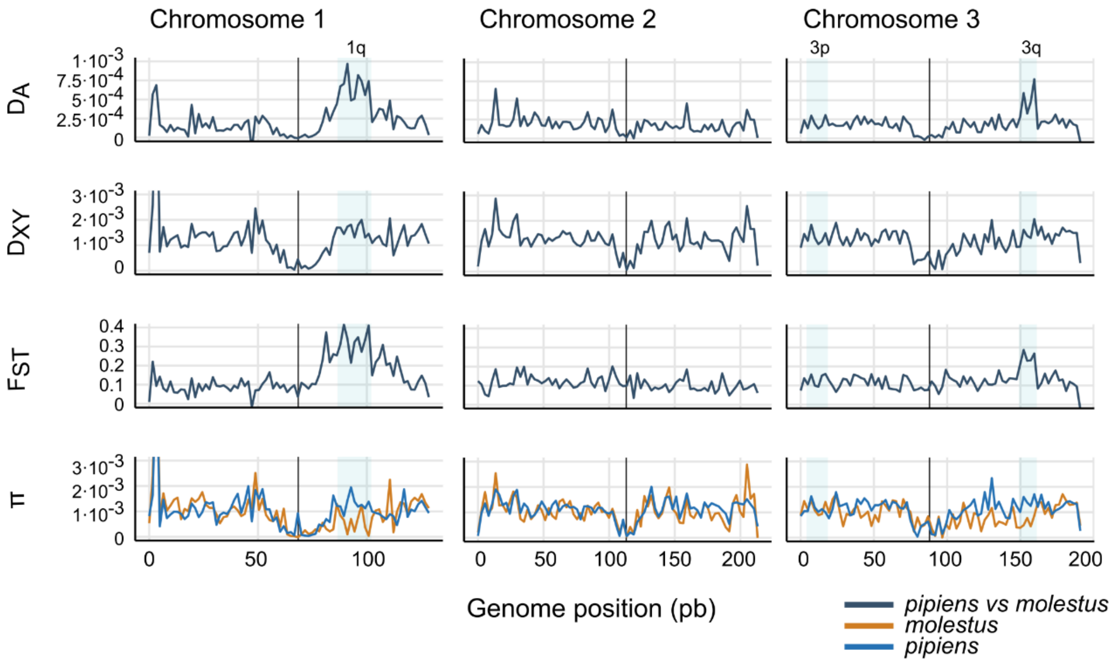


**Supplementary Figure 6: $D_{XY}$, $F_{ST}$ and $D_A$ between *pipiens* and *molestus* and nucleotide diversity (π) estimated over 10kb genomic windows in specimens with 95% or higher ecotype ancestry.** Centromeres are indicated with vertical black lines. Shade blue areas mark putative chromosomal inversions.

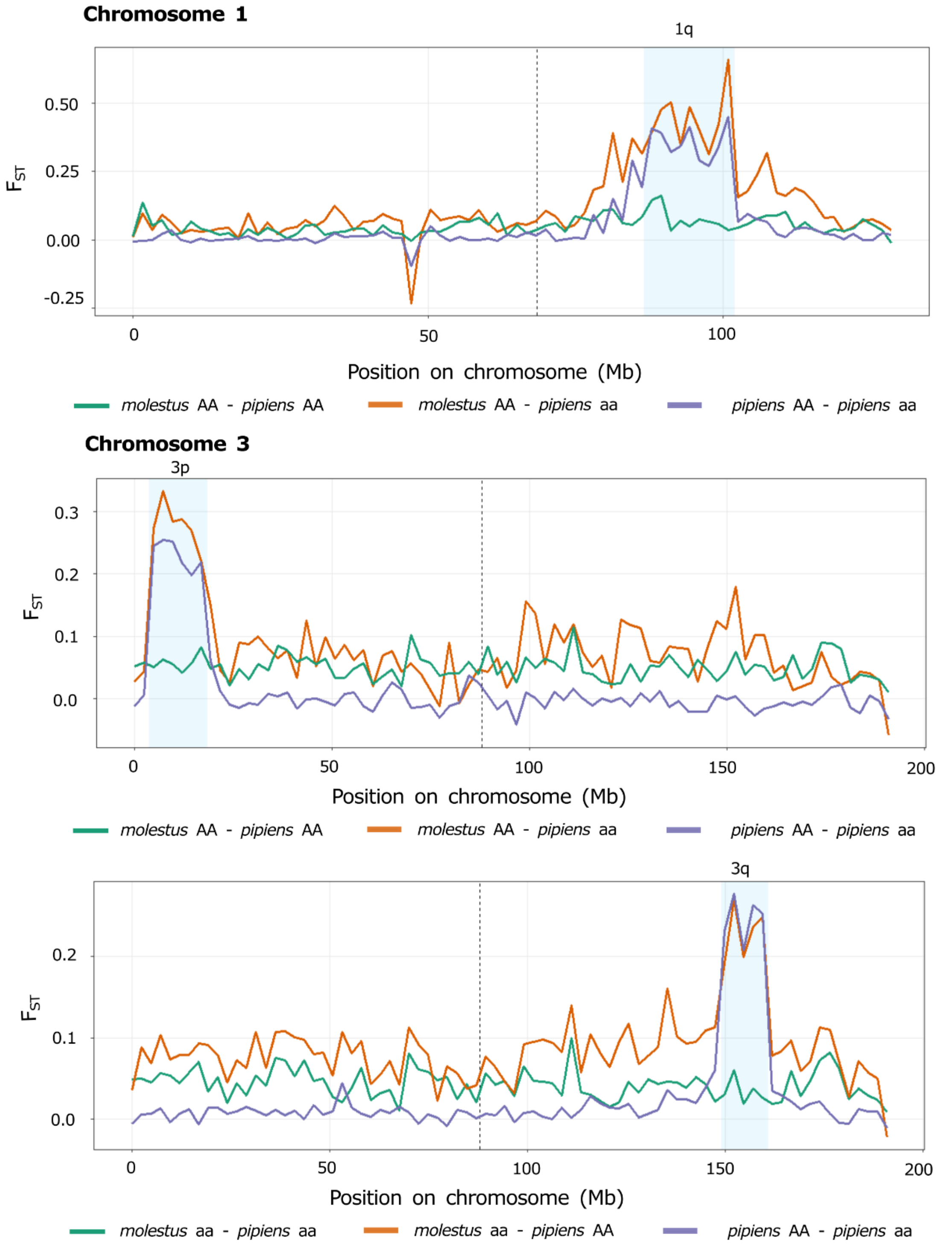


**Supplementary Figure 7: $F_{ST}$ between *pipiens* and *molestus* with standard (AA) and inverted (aa) karyotypes for the three putative chromosomal inversions.** $F_{ST}$ values were estimated over 10kb genomic windows using specimens with 60% or higher ecotype ancestry. Centromeres are indicated with vertical dashed lines. Shade blue areas mark putative chromosomal inversions.

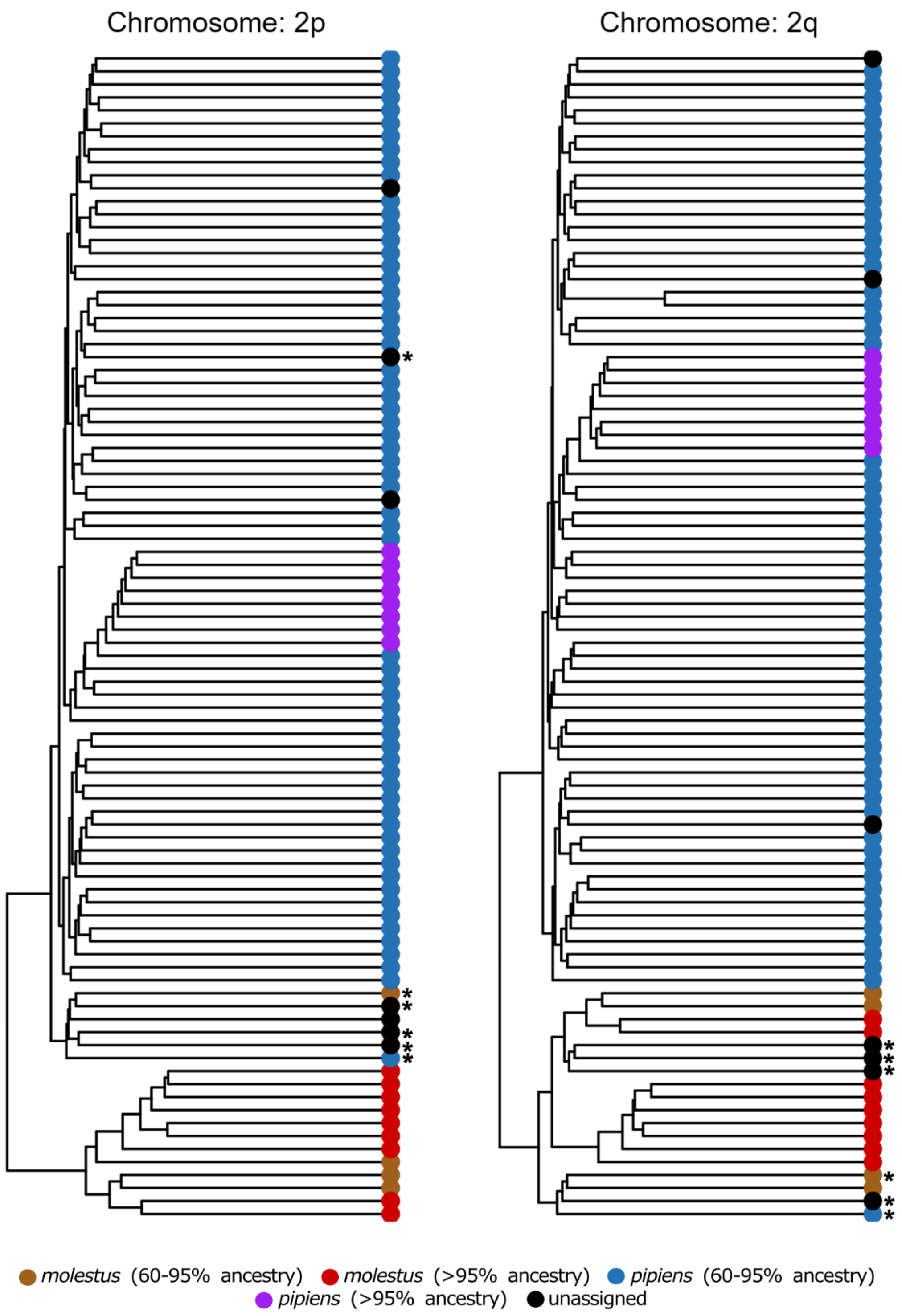


**Supplementary Figure 8:** Phylogenetic reconstruction of 90 *Culex pipiens* based on biallelic SNPs from both arms of chromosome 2. Samples are colored based on ecotype assignment. Samples * indicate samples with discordant lineage assignment between chromosome arms.

### Supplementary Tables

**Supplementary Table 1:** Summary of *Culex pipiens* mosquitoes included in this study. Mosquitoes with >95% ecotype ancestry are shown in parenthesis. Habitat indicates if the sampling location was categorized as urban (U) or natural (N).

| Province | Sampling location | X | Y | *molestus* | *pipiens* | unassigned | N total | Habitat |
|---|---|---|---|---|---|---|---|---|
| Cadiz | Barbate | 36.2302 | -5.8818 | 0 | 2 | 0 | 2 | N |
| | Benalup-Casas Viejas | 36.3567 | -5.7933 | 0 | 9 (4) | 0 | 9 | N |
| | Jerez de la Frontera | 36.614 | -6.0588 | 0 | 9 (5) | 0 | 9 | N |
| | Vejer de la Frontera | 36.2585 | -5.9599 | 0 | 9 | 1 | 10 | U |
| Cordoba | Puente Genil | 37.3808 | -4.7774 | 0 | 10 | 0 | 10 | U |
| Huelva | Corrales | 37.277 | -6.9878 | 6 (6) | 2 | 1 | 9 | U |
| | Huelva | 37.2422 | -6.9274 | 5 | 4 | 0 | 9 | U |
| | La Rabida | 37.2051 | -6.9262 | 0 | 10 | 0 | 10 | U |
| | Los Alamos | 37.2777 | -6.9085 | 2 | 4 | 3 | 9 | U |
| | Moguer | 37.269 | -6.8605 | 0 | 10 | 0 | 10 | U |
| | San Juan del Puerto | 37.3245 | -6.795 | 0 | 9 | 0 | 9 | U |
| Malaga | Campillos | 37.0481 | -4.8342 | 0 | 10 | 0 | 10 | N |
| | Fuente de Piedra | 37.1306 | -4.7424 | 0 | 10 | 0 | 10 | U |
| Seville | Almensilla | 37.2994 | -6.1008 | 1 | 6 | 2 | 9 | U |
| | Bollullos | 37.3151 | -6.1685 | 5 (1) | 5 | 0 | 10 | U |
| | Cañada de los Pajaros | 37.2389 | -6.1289 | 2 | 5 | 2 | 9 | N |
| | Coria del Rio | 37.2828 | -6.0666 | 3 (2) | 5 | 0 | 8 | U |
| | Dehesa de Abajo | 37.2166 | -6.1847 | 0 | 8 (1) | 0 | 8 | N |
| | La Rinconada | 37.477 | -5.9779 | 0 | 6 | 1 | 7 | U |
| | Palomares del Rio | 37.3158 | -6.0566 | 0 | 9 | 1 | 10 | U |
| | Pinares de Colina | 37.2292 | -6.1552 | 0 | 10 (1) | 0 | 10 | U |
| | Puebla del Rio | 37.2596 | -6.074 | 10 (10) | 0 | 0 | 10 | U |
| | San Leandro | 37.0249 | -5.9709 | 0 | 10 | 0 | 10 | N |
| | Seville | 37.4129 | -5.9976 | 0 | 9 (1) | 1 | 10 | U |
| | | | **Total** | 34 (19) | 171 (12) | 12 | 217 | |

**Supplementary Table 2:** Summary of *Culex pipiens pipiens* and *Culex pipiens molestus* samples from Haba *et al*. (2025) included in Principal Component Analyses (PCA) for ecotype assignment confirmation.

| Country | Locality | SRA ID | Sample ID | Sex | Year | *Ecotype* |
|---|---|---|---|---|---|---|
| Belgium | Beveren | SRR31960279 | BVR1 | Female | 2018 | *molestus* |
| | | SRR31960277 | BVR2 | Male | 2018 | *molestus* |
| | | SRR31960276 | BVR3 | Female | 2018 | *molestus* |
| | | SRR32271706 | BVR4 | Male | 2018 | *molestus* |
| | | SRR32271705 | BVR5 | Female | 2018 | *pipiens* |
| | | SRR32271704 | BVR6 | Female | 2018 | *pipiens* |
| | | SRR32271703 | BVR7 | Female | 2018 | *pipiens* |
| | Frameries | SRR31960275; SRR32255677 | FRM1 | Female | 2018 | *pipiens* |
| | | SRR32271711 | FRM2 | Female | 2018 | *pipiens* |
| | | SRR32271710 | FRM3 | Female | 2018 | *pipiens* |
| | | SRR32271709 | FRM4 | Male | 2018 | *pipiens* |
| | Zeebrugge | SRR31960283; SRR32255681 | ZBRb1 | Female | 2017 | *molestus* |
| | | SRR31960281; SRR32255679 | ZBRb3 | Female | 2017 | *molestus* |
| | | SRR31960280; SRR32255678 | ZBRb4 | Female | 2017 | *molestus* |
| | | SRR32271682 | ZBRb5 | Female | 2017 | *molestus* |
| France | Paris | SRR32256194 | PAR1 | Male | 2019 | *pipiens* |
| | | SRR32256193 | PAR2 | Female | 2019 | *pipiens* |
| | | SRR32256158 | PARb1 | Female | 2019 | *molestus* |
| | | SRR32256198 | PARb2 | Male | 2019 | *molestus* |
| | | SRR32256196 | PARb4 | Male | 2019 | *molestus* |
| | | SRR32256195 | PARb5 | Female | 2019 | *pipiens* |
| | | SRR32271964 | PARb6 | Unknown | 2019 | *pipiens* |
| | | SRR32271963 | PARb7 | Unknown | 2019 | *pipiens* |
| Portugal | Setubal | SRR32271840 | BDA1 | Male | 2015 | *pipiens* |
| | | SRR32271839 | BDA2 | Female | 2015 | *molestus* |
| | | SRR32271838 | BDA3 | Female | 2015 | *molestus* |
| | | SRR32271834 | BDA4 | Female | 2015 | *molestus* |
| | | SRR32271842 | BDAi1 | Female | 2015 | *molestus* |
| | | SRR32271841 | BDAi2 | Female | 2015 | *pipiens* |
| | | SRR32271836 | BDAi3 | Female | 2015 | *pipiens* |
| | | SRR32271835 | BDAi4 | Female | 2015 | *molestus* |
| | | SRR32271923 | BDAi5 | Female | 2015 | *molestus* |
| | | SRR32271922 | BDAi6 | Female | 2015 | *molestus* |
| Spain | Extremadura | SRR32271823 | EXT1 | Female | 2018 | *pipiens* |
| | Garrapinillos | SRR32280606 | GPN1 | Female | 2019 | *pipiens* |
| | | SRR32280605 | GPN2 | Female | 2019 | *pipiens* |

| | | | | | |
|---|---|---|---|---|---|
| | SRR32280604 | GPN3 | Female | 2019 | *molestus* |
| | SRR32271828 | GPN4 | Female | 2019 | *pipiens* |
| | SRR32271827 | GPN5 | Female | 2019 | *pipiens* |
| Zaragoza | SRR32269686 | GPNi1 | Female | 2019 | *molestus* |
| | SRR32269685 | GPNi2 | Female | 2019 | *molestus* |
| | SRR32271825 | GPNi4 | Female | 2017 | *pipiens* |
| | SRR32271824 | GPNi5 | Female | 2017 | *molestus* |
| Seville | SRR32271833 | MDA1 | Female | 2019 | *pipiens* |
| | SRR32271832 | MDA2 | Female | 2019 | *pipiens* |
| | SRR32271831 | MDA3 | Female | 2019 | *pipiens* |
| | SRR32271830 | MDA4 | Female | 2019 | *pipiens* |
| | SRR32271829 | MDA5 | Female | 2019 | *pipiens* |

**Supplementary Table 3**: Karyotype frequencies estimated for three putative chromosomal inversions over 217 *Culex pipiens*. Mosquitoes with >95% ecotype ancestry are indicated in brackets. NA corresponds to mosquitoes in which karyotype assignment was not possible.

| Inversion | Karyotype | *molestus* | *pipiens* | unassigned | Total |
|---|---|---|---|---|---|
| 1q | wildtype | 31 (19) | 104 | 11 | 146 |
| | heterokaryotype | 2 | 54 (6) | 1 | 57 |
| | inverted | | 11 (6) | | 11 |
| | NA | 1 | 2 | | 3 |
| 3p | wildtype | 32 (19) | 109 (12) | 11 | 152 |
| | heterokaryotype | 2 | 54 | 1 | 57 |
| | inverted | | 7 | | 7 |
| | NA | | 1 | | |
| 3q | wildtype | | 21 (12) | 1 | 22 |
| | heterokaryotype | | 63 | 4 | 67 |
| | inverted | 34 (19) | 87 | 7 | 128 |

**Supplementary Table 4:** Gene Ontology (GO) enrichment analysis restricted to chromosomal inversions. Only GO terms significant at α = 0.05 after false discovery rate (FDR) correction are shown.

| Inversion | GO Term | Function | Category | Count | Total genes | p-value | FDR |
|---|---|---|---|---|---|---|---|
| 3p | GO:0050911 | detection of chemical stimulus involved in sensory perception of smell | BP | 14 | 103 | 1.70E-07 | 7.06E-04 |
| | GO:0050896 | response to stimulus | BP | 53 | 1344 | 2.60E-07 | 7.06E-04 |
| | GO:0140359 | ABC-type transporter activity | MF | 14 | 55 | 2.90E-10 | 5.77E-07 |
| | GO:0020037 | heme binding | MF | 20 | 181 | 2.30E-07 | 2.29E-04 |
| | GO:0004984 | olfactory receptor activity | MF | 14 | 103 | 1.30E-06 | 8.61E-04 |
| | GO:0005506 | iron ion binding | MF | 19 | 199 | 4.30E-06 | 2.14E-03 |
| | GO:0016705 | oxidoreductase activity, acting on paired donors, with incorporation or reduction of molecular oxygen | MF | 19 | 205 | 6.70E-06 | 2.39E-03 |
| | GO:0004497 | monooxygenase activity | MF | 19 | 206 | 7.20E-06 | 2.39E-03 |
| | GO:0016887 | ATP hydrolysis activity | MF | 20 | 229 | 9.40E-06 | 2.67E-03 |
| | GO:0005789 | endoplasmic reticulum membrane | CC | 23 | 295 | 1.00E-05 | 9.39E-03 |
| 3q | GO:0017121 | plasma membrane phospholipid scrambling | BP | 5 | 7 | 3.90E-08 | 2.12E-04 |
| | GO:0042302 | structural constituent of cuticle | MF | 16 | 123 | 7.80E-09 | 1.55E-05 |
| | GO:0016747 | acyltransferase activity, transferring groups other than amino-acyl groups | MF | 16 | 158 | 6.90E-08 | 6.03E-05 |
| | GO:0017128 | phospholipid scramblase activity | MF | 5 | 7 | 9.10E-08 | 6.03E-05 |
| | GO:0005212 | structural constituent of eye lens | MF | 3 | 3 | 9.90E-06 | 4.92E-03 |

| | | | | | | | |
|---|---|---|---|---|---|---|---|
| | GO:0008609 | alkylglycerone-phosphate synthase activity | MF | 3 | 5 | 9.60E-05 | 3.82E-02 |
| | GO:0045277 | respiratory chain complex IV | CC | 3 | 4 | 1.80E-05 | 1.69E-02 |
| 1q | GO:0006520 | amino acid metabolic process | BP | 10 | 153 | 5.00E-07 | 2.71E-03 |

**Supplementary Table 5**: f3 statistics for three different combinations of source and target populations between *Culex pipiens pipiens*, *Culex pipiens molestus* and samples of *Culex pipiens* unassigned to any of the ecotypes. Significant results indicating signs of admixture are shown in bold (|Z| ≥ 3).

| Target | Source1 | Source2 | f_3 | std.err | Z |
|---|---|---|---|---|---|
| unassigned (N=7) | *molestus* | *pipiens* | -9.32E-04 | 3.78E-04 | -2.467 |
| molestus (N=13) | *pipiens* | unassigned | 6.01E-04 | 2.19E-04 | 2.746 |
| **pipiens (N=70)** | **unassigned** | ***molestus*** | **-5.83E-04** | **1.68E-04** | **-3.46** |

**Supplementary Table 6**: Reference *pk1* sequences from different *w*Pip groups used to classify *w*Pip consensus assemblies from 217 *Culex pipiens.*

| Accession number | ***Strain*** | Reference |
|---|---|---|
| OM885005.1 | wPip-I | (Atyame *at al.*, 2011) |
| OM885006.1 | wPip-III | (Atyame *at al.*, 2011) |
| OM885007.1 | wPip-II | (Atyame *at al.*, 2011) |
| OM885008.1 | wPip-IV | (Atyame *at al.*, 2011) |
| OM885009.1 | wPip-V | (Atyame *at al.*, 2011) |
| LC106155.1 | wPip-I | (de Pinho Mixão *et al*., 2016) |
| OQ223315.1 | wPip-II | (da Moura *et al*., 2023) |
| OQ223310.1 | wPip-III | (da Moura *et al*., 2023) |
| OQ223309.1 | wPip-IV | (da Moura *et al*., 2023) |
